# GTP cycling gates CTP synthase activation

**DOI:** 10.1101/2022.09.24.509316

**Authors:** Chen-Jun Guo, Yu-Fen Wu, Shu-Ying Guo, Jia-Li Lu, Liang Xu, You Fu, Xian Zhou, Jiale Zhong, Wei Wang, Zherong Zhang, Ji-Long Liu

## Abstract

CTP synthase (CTPS), the sole enzyme for de novo CTP biosynthesis, requires GTP as an allosteric effector for efficient catalysis^1^. Although the labile intermediates and chemical reaction steps have been characterized, how CTPS coordinates its structural domains through interactions with GTP to continuously catalyze thus reactions and thereby synthesize CTP correctly remains elusive. Through cryo-EM analysis of 34 structural states (2.0–3.3 Å resolution) spanning pre-catalytic, catalytic, and post-catalytic conformations, we delineate how GTP cycling—intact binding and dissociation—directly gates three sequential activation checkpoints. 1) Catalytic initiation gate: GTP dissociation precedes substrate entry, transitioning CTPS to a substrate-competent state. 2) Intermediate-dependent GTP recruitment: 4Pi-UTP formation allosterically remodels the GAT domain, licensing GTP rebinding. 3) Ammonia transfer gate: GTP directly constitutes the transient interdomain gas tunnel for ammonia delivery while stimulating the glutamine hydrolyzing. CTP synthesis completion triggers GTP dissociation, allowing glutamate release. Each catalytic cycle couples CTP production to a single GTP-binding event, demonstrating that GTP cycling operates as a catalytic-phase timer. This work provides a comprehensive model for the efficient catalysis of CTPS and establishes effector cycling as a dynamic gating mechanism in enzymes.

## Main

CTP serves as an essential precursor for RNA synthesis and phospholipid biosynthesis, with its intracellular levels tightly regulated to maintain nucleotide homeostasis^2-4^. In living organisms, CTP is synthesized de novo by the enzyme CTP synthase (CTPS). Given the central metabolic role of CTP, CTPS has emerged as a potential therapeutic target for autoimmune diseases, cancer, and infections caused by viruses, prokaryotes, and protozoa^5-11^. Additionally, recent studies have revealed that CTPS possesses the catalytic ability to deamidate protein and plays important roles in antiviral innate immunity and DNA repair processes^12,13^.

CTPS is a multifunctional enzyme composed of two functional domains: an N-terminal Amidoligase (AL) domain and a C-terminal Glutamine Amide Transferase (GAT) domain^14,15^ **(Fig. 1a)**. CTPS typically utilizes glutamine as a nitrogen donor and catalyzes the synthesis of CTP from UTP through a three-step reaction mechanism: (1) The AL domain catalyzes the formation of an unstable intermediate, 4Pi-UTP, consuming ATP and producing ADP. (2) The GAT domain hydrolyzes glutamine to generate ammonia, which is channeled through a transient internal tunnel to the AL domain. (3) The ammonia then reacts with 4Pi-UTP in the AL domain to produce the final product, CTP^16-18^ **(Fig. 1b)**.

**Figure 1.**
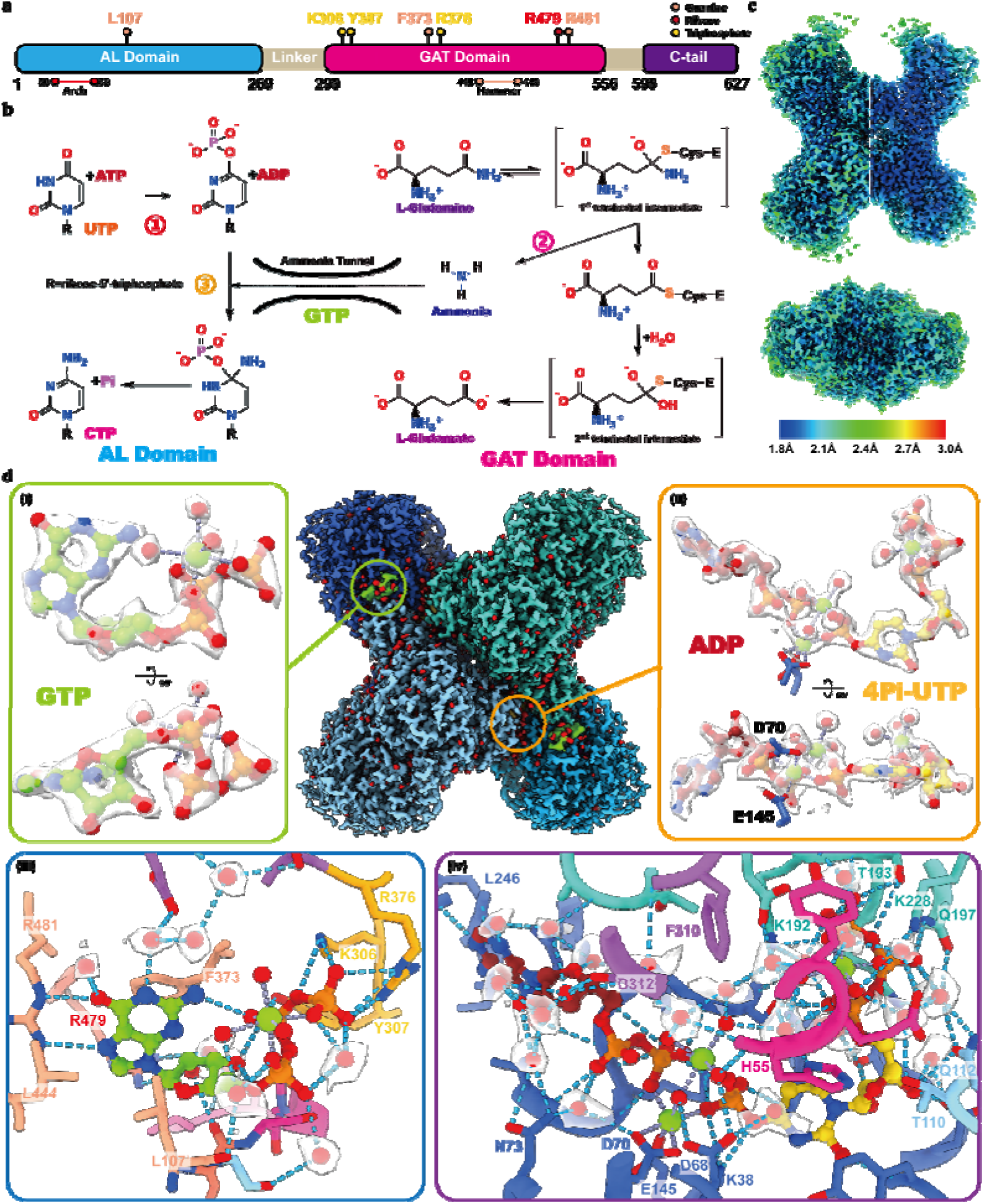
Structure of the CTPS with ADP, 4Pi-UTP, GTP and DON at 2.0 Å resolution. **a**, Domain organization of CTPS with GTP-binding residues color-coded by interacting functional groups. **b**, Reaction catalyzed by CTPS, coupling catalytic cycles with ammonia tunneling. Reaction 1 occurs in the AL domain, where ATP and UTP are consumed to produce ADP and 4Pi-UTP. The GAT domain hydrolyzes glutamine to generate ammonia via the formation and collapse of two tetrahedral intermediates. The intermediate product, ammonia, is transferred from the GAT domain to the AL domain through a transient internal gas tunnel composed of both protein and the GTP molecule. Finally, 4Pi-UTP is aminated by ammonia to produce the end product, CTP. **c**, Local resolution map of the 2.0 Å CTPS structure. **d**, Visualization of chemical details in the 2.0 Å structure, enabling precise assignment of Mg²L coordination geometry. The CTPS tetramer is colored by protomer. Water molecules are shown in red. The locations of zoomed-in views (i) and (ii) are indicated. (i) and (ii): Map density and atomic models of GTP or ADP and 4Pi-UTP, coordinated with magnesium ions and water molecules. (iii) and (iv): Hydrogen bonding networks stabilising allosteric effector and catalytic intermediates (iii: GTP pocket; iv: ADP and 4Pi-UTP pocket). Densities corresponding to surrounding water molecules are shown. The models are colored according to the scheme presented in panel **a**.

GTP is essential for the efficient catalysis by CTPS and serves as a critical regulator of its activity^19^. The recognition of GTP by CTPS is highly specific^20^. Although GTP is not consumed during the reaction, its effect is dose-dependent and multifaceted: it can either activate or inhibit CTPS activity depending on the context^21,22^. Owing to the paucity of structural insights, the precise role of GTP in CTPS regulation has remained elusive, akin to a spectral presence whose influence is inferred yet not fully elucidated. In our previous work, using an irreversible inhibitor, we captured the first structure of CTPS bound to GTP, confirming their direct interaction^23^. However, this structure revealed a paradox: while GTP binding enabled the transfer of reaction intermediates, it simultaneously appeared to block the catalytic pocket preventing product release—effectively “locking” the enzyme. This raised a critical question: does this conformation exist under physiological ligand conditions? How exactly does GTP modulate and promote CTPS activity?

In this study, we employed a reaction-based time-resolved cryo-EM strategy combined with continuous heterogeneity analysis to systematically resolve a series of conformational states of CTPS under native ligand conditions—before the reaction, during catalysis, and at equilibrium^24^. These structural snapshots reveal the dynamic interplay between CTPS and GTP throughout the catalytic cycle, from substrate binding and catalysis to product formation. Cross-validation using three different inhibitors conditions enabled us to identify key ‘checkpoint’ conformations, including a 2.0 Å resolution structure that allows visualization of delicate intermediates and solvent molecules. In total, 34 high-quality structures representing distinct states of CTPS are obtained, presenting a comprehensive, empirically derived mechanism for GTP-driven CTP synthesis that may be conserved across all domains of life. As such, we demonstrate a novel allosteric regulation mechanism that effector allosteric regulates enzyme by the cyclic binding during the reaction cycle.

## Results

### Structure of CTPS with GTP and intermediates at 2 Å resolution

To obtain a high-resolution structure of CTPS in its GTP-bound state, we employed the irreversible covalent inhibitor 6-Diazo-5-oxo-L-norleucine (DON) to substitute for the substrate glutamine, thereby stabilizing the sample. Cryo-EM grids were prepared in the presence of UTP, ATP, GTP, and Mg^2+^, referred to as the DON-state. Following data acquisition on a 300 kV Titan Krios microscope, we processed and reconstructed the data, yielding an electron density map at a nominal resolution of 2.0 Å **(Fig. S1)**. The local resolution of the map varied from 1.8–2.0 Å at the AL domain and 1.9–2.7 Å at the GAT domain **(Fig. 1c)**. This resolution enabled confident modeling of amino acid side chains and revealed interactions between metal ions, ordered solvent molecules, and ligands **(Fig. 1d, S2 and Supplementary Videos 1 and 2)**.

In the GTP-binding site, we observed a magnesium ion directly coordinated to the GTP molecule **(Fig. 1d)**. This ion adopts an octahedral geometry, forming coordination bonds with one oxygen atom each from the α, β, and γ phosphates of GTP, with the remaining three positions occupied by water molecules. Within the AL domain, we identified three magnesium ions interacting with the reaction intermediate 4Pi-UTP and ADP, exhibiting octahedral coordination **(Fig. 1d)**. One magnesium ion stabilizes the three phosphate groups of 4Pi-UTP by coordinating with one oxygen from each phosphate group and three water molecules. At the junction between 4Pi-UTP and ADP, we observed two additional magnesium ions: one bridges the β-phosphate of ADP, the 4-position phosphate of 4Pi-UTP, and four water molecules. The second bridges the β-phosphate of ADP, the 4-position phosphate of 4Pi-UTP, and carboxylate side chains from E145 and D70, as well as one water molecule **(Fig. S2b)**.

GTP is stabilised by multiple interactions **(Fig. 1d, S3a-d and Supplementary Video 1)**: it adopts a folded-back conformation supported by intramolecular hydrogen bonds between N2 and the β-phosphate, as well as between the O3′ of the ribose and the α/β phosphates. The triphosphate moiety is recognized by K306, Y307, and R376 through multiple hydrogen bonds and electrostatic interactions, with two water molecules contributing additional stabilization—one of which mediates a contact with G108′. The ribose is anchored via hydrogen bonds with F50 and R479. The guanine base forms two hydrogen bonds with R481, engages in π– π stacking with F373, and contacts with L107. A water molecule mediates its contact with E423. These extensive interactions explain the high specificity of CTPS for GTP^25,26^.

Sequence comparisons across species from different domains of archaea, bacteria, and eukaryote revealed that the residues involved in GTP binding are highly conserved, confirming GTP cycling as an ancient regulatory strategy **(Fig. S3e-h)**.

In AL domain, a hydrogen-bond network involving both sidechain and ordered solvent molecules stabilises the delicate intermediate 4Pi-UTP **(Fig. 1d and Supplementary Video 2)**. The 50–58 loop, which constitutes the core of the ammonia tunnel, contains residues P52, E57, and V58—previously proposed to act as the “bottleneck” of the tunnel, hereafter referred to as the Arch^23,27-29^. The Arch has been implicated in UTP recognition and conformational transitions that promote tunnel formation, though the precise mechanism has remained unclear. We found that the Arch recognizes 4Pi-UTP through multiple bridging hydrogen bonds mediated by ordered solvent: the side chain of H55 interacts with both the triphosphate of 4Pi-UTP and a coordinating magnesium ion via a water molecule; the E54 side chain forms hydrogen bonds with the triphosphate through two distinct ordered water molecules.

### Capturing the dynamic process of GTP and CTPS interaction

In the DON-state structure, GTP forms a tight binding with CTPS. This interaction stabilises a closed conformation essential for ammonia tunnel integrity **(Fig. S4a, b)**. However, this static snapshot paradoxically blocks product release (**Fig. S4c and Supplementary Video 3**), necessitating dynamic cycling for sustained catalysis.

To visualize this intricate mechanism, we established a reaction-based time-resolved cryo-EM system to track GTP behaviour across catalytic stages: 1. Pre-reaction (PRE): CTPS with ATP/UTP/GTP, no glutamine; 2. Active reaction (RXN): Post-glutamine addition (0–180 s); 3. Equilibrium (EQ): CTP-accumulated inhibition (>900 s) **(Fig. S5a)**. Complementary inhibitor strategies isolated specific steps: 4. ACP-state (AMP-PCP substitution) arrested AL-domain activity; 5. dDON-state (DON+dNTPs) inhibited the GAT-domain and slowed the catalysis^30^.

Following data acquisition and initial reconstructions, we observed structural heterogeneity under each condition, indicating the presence of multiple conformational states. To resolve these dynamics in greater detail, we focused on a single CTPS protomer and performed focused 3D classification. First, particles from homogeneous refinement were symmetry-expanded and subjected to 3D classification using a focused mask on one protomer, with the resolution restricted at 6 Å to highlight large conformational differences. Damaged or misaligned particles were removed based on this low-resolution classification. Next, we applied 3D variability analysis (3DVA) to dissect conformational changes further. 3DVA decomposes structural heterogeneity into a series of principal components ranked by their contribution to overall variance. Principal component 0 (PC0), the most dominant mode, was used to cluster particles using a window size of 0, yielding 4 to 8 non-redundant conformational classes. Individual reconstructions of each cluster revealed a range of discrete intermediate states, enabling detailed structural analysis **(Fig. 2a and Supplementary Video 4-13)**.

**Figure 2.**
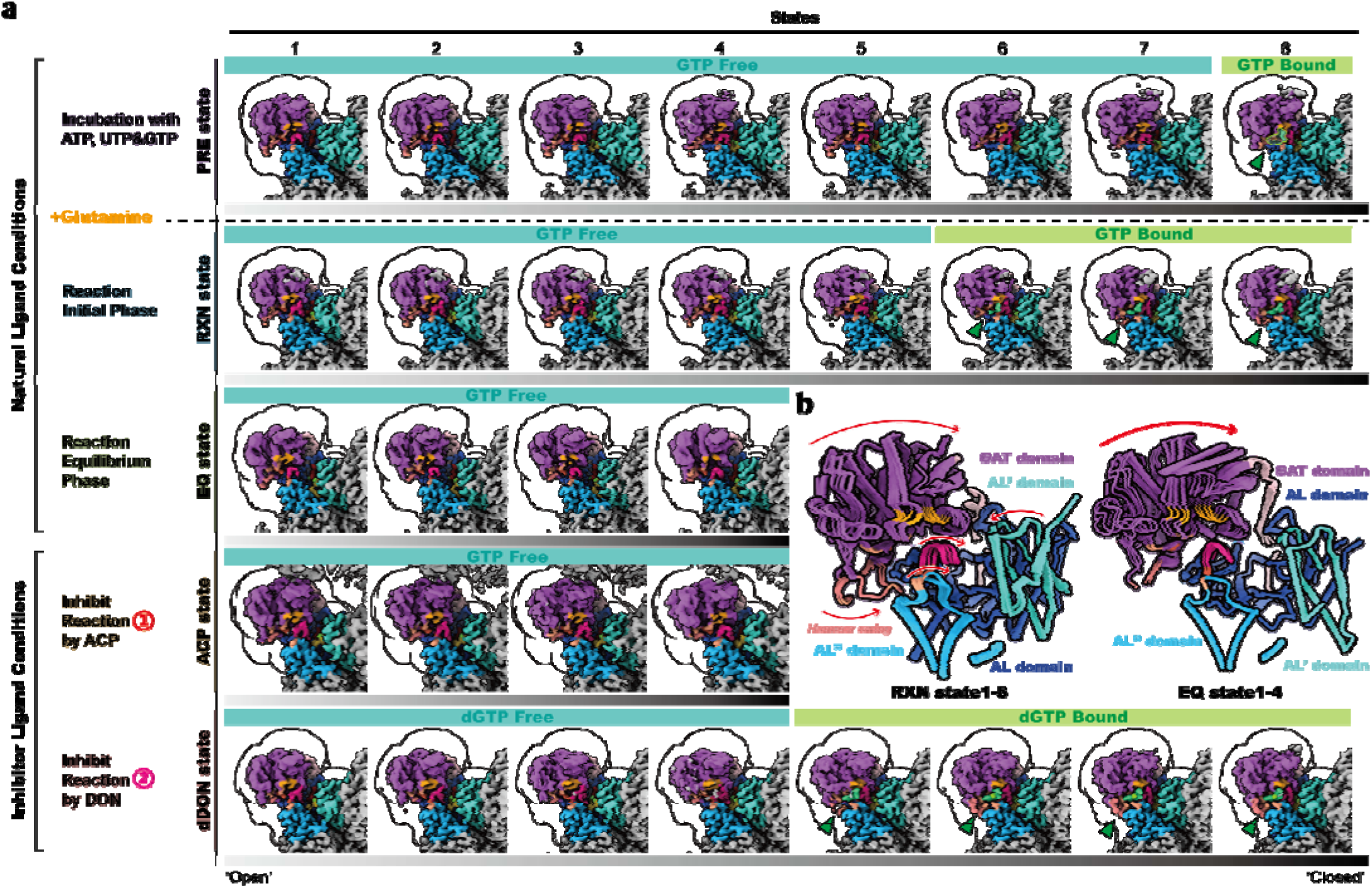
GTP binding gates catalytic progression through conformational selection. **a**, CTPS structural ensembles under reaction conditions. Conformational dynamics of CTPS revealed by cryo-EM. Maps are grouped according to sample preparation conditions. One protomer and two AL domains from adjacent protomers are shown, with different colors distinguishing each component. The mask used for focused classification is represented as a transparent white surface. States with GTP binding are indicated. Color bars beneath each structural ensemble illustrate the prevalence of the ‘open’ or ‘closed’ conformation within each condition. All maps are displayed at the same contour level=0.1 for consistent comparison. **b**, Model comparison of structural ensembles from two natural ligand condition. The left panel shows the RXN-state, while the right panel displays the EQ-state.

This workflow was applied to datasets from five CTPS sample conditions, yielding a set of intermediate structures at 2.8–3.3 Å global resolution **(Fig. S6-10)**. In the PRE-state, dDON-state, and RXN-state, we captured snapshots of GTP binding, revealing GTP cycling synchronises CTPS catalytic transitions. In contrast, no GTP binding was observed in the EQ- or ACP-states. Across all datasets, CTPS exhibited relative movements between the GAT and AL domains. We refer to conformations with greater interdomain distance as the open conformation, and those with shorter distances as the closed conformation **(Fig. 2b)**.

### GTP binding requires AL domain catalysis

In the PRE-state sample, we capture the formation of 4Pi-UTP **(Fig. 3a)**. In the electron density map of state 1, ATP is bound and stabilized, with the γ-phosphate connected to the β-phosphate, indicating that reaction 1 has not yet occurred. At the UTP-binding site, only weak density is observed near the triphosphate moiety. By state 3, density corresponding to the UTP head group connects to the γ-phosphate of ATP to form 4Pi-UTP, while the β- and γ-phosphates become separated—indicating that the first step of catalysis in the AL domain has taken place. In states 6 and 8, we observed well-resolved densities corresponding to ADP and 4Pi-UTP, as well as a shift of the Arch motif toward the AL domain. Focusing on the GTP-binding pocket, we found that GTP binding was observed only after reaction 1 had occurred within the AL domain. Specifically, no GTP density was detected in states 1, 3, or 6, but GTP is clearly resolved in state 8. Comparing state 1 with state 8, the formation of 4Pi-UTP is accompanied by the inward movement of the Arch and GAT domains, with the Hammer region becoming increasingly disordered in the process **(Fig. 3b)**.

**Figure 3.**
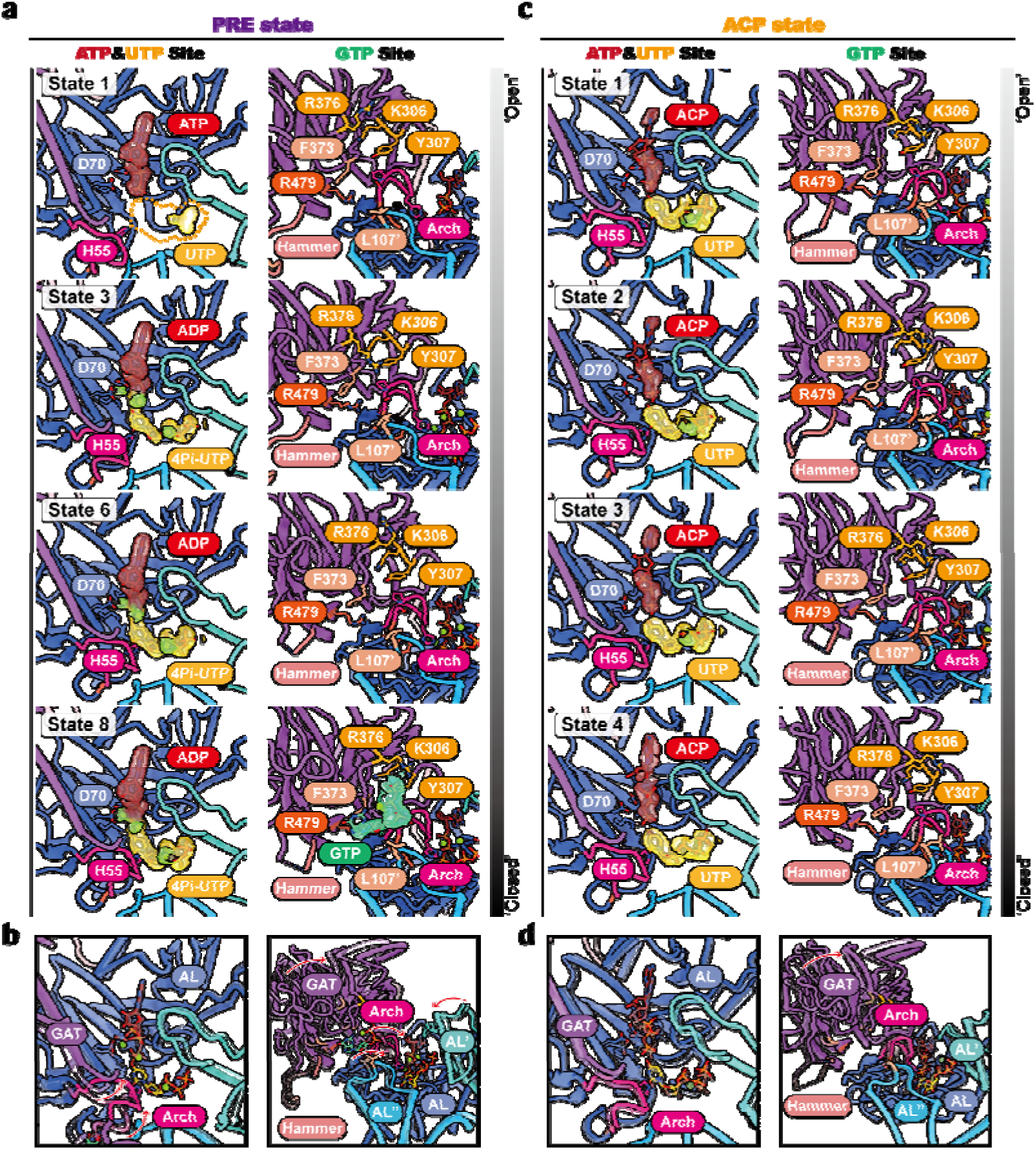
AL domain catalysis licenses GTP binding through structural proofreading. **a**, Cryo-EM structural ensemble from PRE state showing progressive AL domain closure (stages 1-6) proceeds GTP binding (stage 8). The ligand-binding states of the AL domain and GTP-binding pocket are shown. Ligand densities are displayed as transparent surfaces. **b**, Structural comparison between state 1 and state 8 from the PRE-state sample. The comparison focuses on the AL domain (left) and the overall conformational changes (right). **c**, ACP-state structures demonstrate failed AL-GAT coupling despite AL domain movement (left) and zero GTP density across all states (right). The ligand-binding states of the AL domain and GTP-binding pocket are shown. Ligand densities are displayed as transparent surfaces. **d**, Structural comparison between state 1 and state 8 from the ACP-state sample. The comparison focuses on the AL domain (left) and the overall conformational changes (right).

GTP exhibits weak inhibition of NH -dependent activity in *E. coli* CTPS, indicating that GTP can bind in the presence of ATP and UTP^21,31^. Our structural findings support this conclusion and provide a mechanistic basis. We further validated this using *Drosophila melanogaster* CTPS, where we observed similar inhibitory effects at high GTP concentrations under NH -dependent conditions **(Fig. S5b)**. These data show that GTP can bind CTPS in the absence of GAT domain ligands. Given that states 1 through 8 were derived from the same dataset, GTP binding is conformation selective, with the formation of 4Pi-UTP serving as a prerequisite—a checkpoint—for GTP association.

To test whether catalysis in the AL domain is required for GTP binding, and whether ligand binding alone is sufficient to induce GTP association, we analyzed the ACP-state sample, in which AL domain activity is blocked by replacing ATP with the non-hydrolyzable analog ACP.

In the ACP-state, although we observed movement of the GAT domain toward the AL domain, no stable GTP binding was detected **(Fig. 3c, d)**. Zooming in on the AL domain, the three phosphate groups of ACP remained intact in all ACP-state reconstructions, and no formation of 4Pi-UTP was observed—indicating that reaction 1 was successfully inhibited. At the GTP-binding pocket, no GTP density was observed in any of the four states.

These findings collectively demonstrate that substrate binding and associated conformational changes in the AL domain are insufficient for GTP binding. Instead, the occurrence of reaction 1 and formation of 4Pi-UTP is essential, positioning it as a structural checkpoint for GTP binding.

### GTP binding is further stabilized by ligand-bound GAT domain

Analysis of the dDON-state dataset revealed the transition of CTPS from a GTP-free to a GTP-bound state, along with structural insights into how the ligand-bound GAT domain promotes GTP stabilization **(Fig. 4a)**. In all reconstructed states, we observed clear density for DON at the GAT pocket, confirming its occupancy. In state 1, both dATP and dUTP were intact, and no density for dGTP was detected, indicating that reaction 1 had not occurred. In state 3, formation of dADP and 4Pi-dUTP marked the completion of reaction 1, along with a shift of the Arch and GAT domains toward AL. dGTP remained unbound. In state 5, further movement of Arch and GAT toward AL was observed, and dGTP binding emerged. Here, R481 reoriented toward dGTP, and L107 adopted a new conformation to support the guanine base. The Hammer (Loop^440-448^) underwent a conformational swing from the exterior of GAT toward the AL domain. In state 8, R481 moved closer to dGTP, establishing two hydrogen bond interactions, and the Hammer density was further stabilized, indicating a fully engaged GTP-binding mode. Compared to PRE-state 8, states dDON-5 and dDON-8, with ligand-bound GAT domains, showed significantly enhanced GTP stabilization.

**Figure 4.**
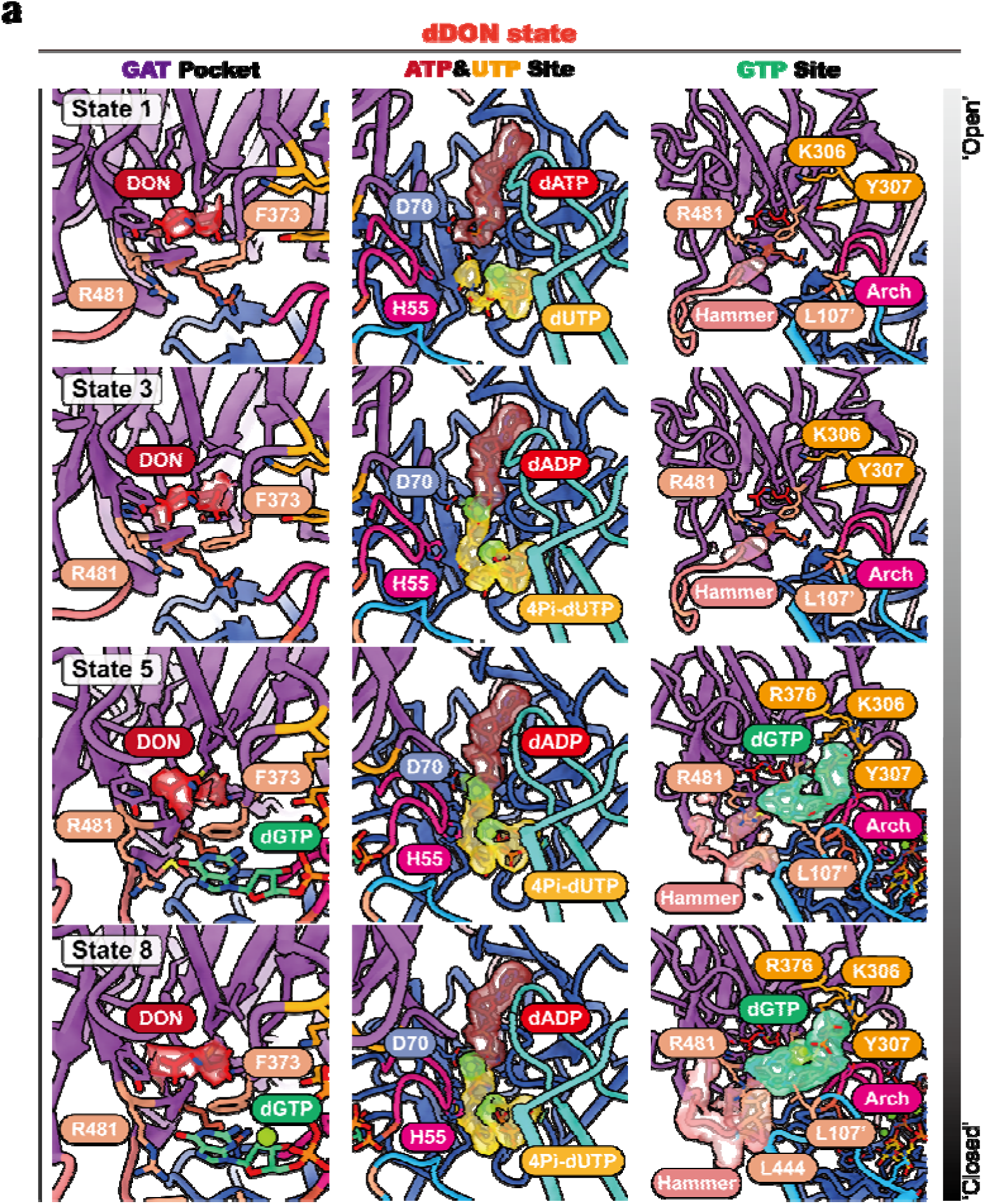
Ligand-bound GAT domain mediates dGTP stabilization in dDON sample. **a**, Models of selected states from the dDON-state sample shows the allosteric synergy between dGTP and inhibitor DON. The ligand-binding states of the GAT domain, AL domain and GTP-binding pocket are shown. Ligand and hammer densities are displayed as transparent surfaces.

We observed similar patterns in the natural-ligand RXN-state dataset **(Fig. 5a)**: In state 1, ATP and UTP remained intact, and GTP was unbound. In state 3, reaction 1 occurred, marked by ADP and 4Pi-UTP formation, and GAT and Arch shifted toward AL, but GTP did not bind. In state 6, further movement of GAT and Arch coincided with GTP binding, R481 reorientation, and conformational changes in the hammer. In state 8, L107 flipped to support GTP, R481 formed two hydrogen bonds with the guanine base, and the Hammer became fully stabilized by folding toward the AL domain. Moreover, our biochemical assays indicate that GTP binding promotes GAT domain activity, consistent with previous observations in ecCTPS and llCTPS^21,22^ **(Fig. S5c)**.

**Figure 5.**
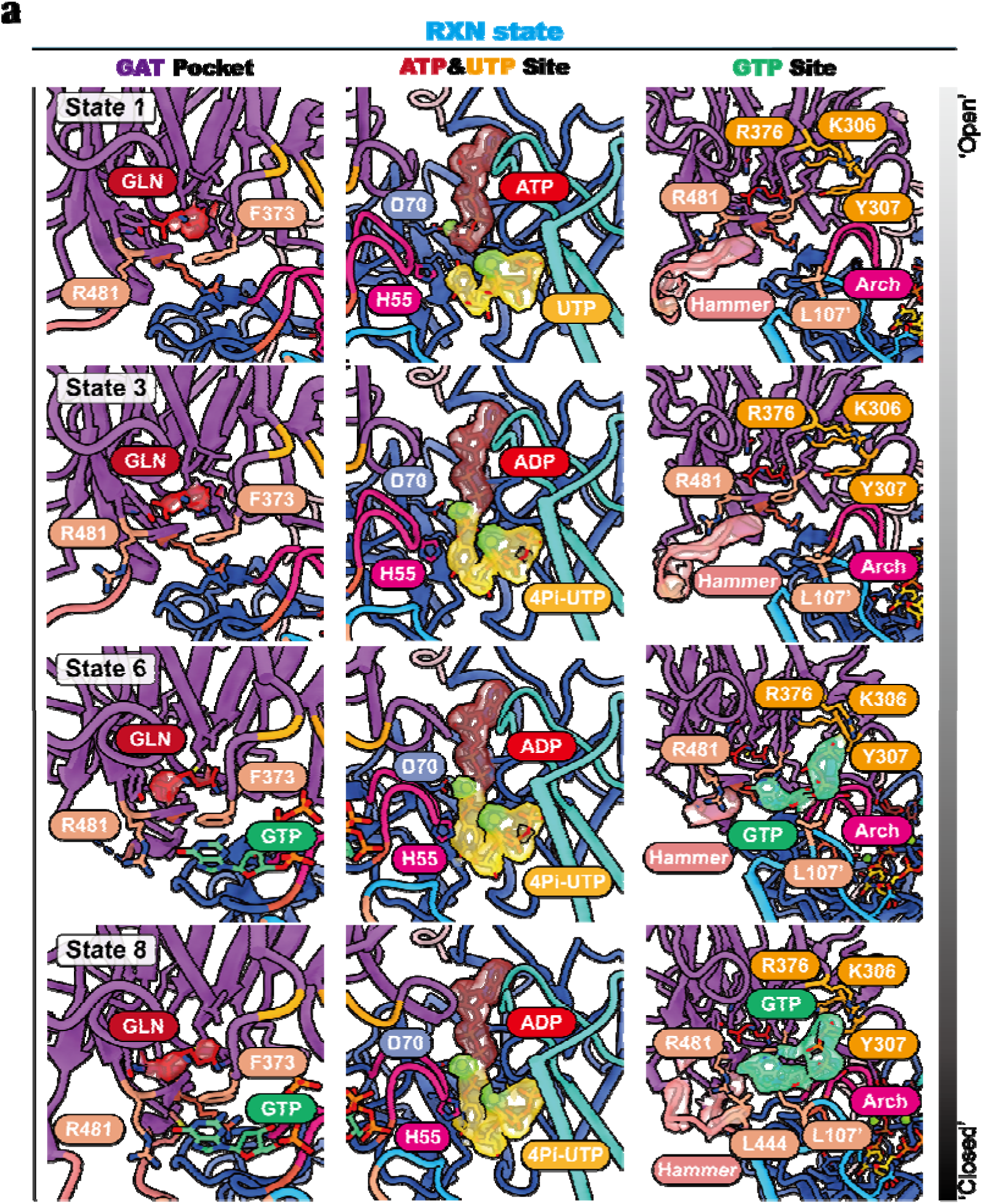
Ligand-bound GAT domain mediates GTP stabilization in RXN sample. **a**, Models of selected states from the RXN-state sample. The ligand-binding states of the GAT domain, AL domain and GTP-binding pocket are shown. Ligand and hammer densities are displayed as transparent surfaces.

Together, our findings from both the dDON-state and RXN-state indicate that ligand binding in the GAT domain promotes GTP stabilization. Beyond enhancing enzymatic activity, GTP binding also contributes to closing the ammonia channel, securing the GAT pocket, preventing substrate dissociation, and blocking additional ligand competition—underscoring its role as both a regulatory and structural gatekeeper in the catalytic cycle.

### The side-chain volume of L444 is critical for CTPS catalysis

In the DON-state, which mimics the GAT domain in a glutamine-binding or tetrahedral intermediate formation state, L444 is located near the GTP binding site and contacts with R481 and L107 **(Fig. S11a)**. The amino acids interacting with GTP in CTPS are highly conserved. Multiple sequence alignment revealed diversity at position L444, including methionine and lysine **(Fig. S11b-c)**. Our previous study showed that mutating L444 to alanine abolishes CTPS activity^23^. To further probe its role, we performed site-directed mutagenesis and found that substitutions with methionine, isoleucine, phenylalanine, and lysine—retained enzymatic activity. In contrast, valine, which has a smaller side chain than leucine, exhibited reduced activity, while threonine substitutions led to the loss of function **(Fig. S11d)**. Structural analyses of both RXN-state and dDON-state datasets revealed conformations in which GTP is stably bound but the hammer density is unresolved. The conformational change of the Hammer region consistently occurs after the interactions between GTP and other amino acids. Taken together with the biochemical data, we propose that, unlike other residues in the GTP pocket, L444 does not inherently stabilize GTP binding. Instead, it interacts with GTP and surrounding amino acids through the inward movement of the Hammer, generating steric hindrance that triggers the collapse of the tetrahedral intermediate formed during the first reaction in the GAT domain— functioning like a firing pin to ultimately promote ammonia release.

### GTP unbinds post-reaction

During 3D classification of the RXN-state dataset, we identified a small portion of particles in a CTP-bound state, referred to as the p-state **(Fig. 6a)**. In the RXN p-state structure, GTP binding was not observed. ATP occupies in the ATP-binding pocket, while CTP occupies the UTP pocket. This CTP-binding mode is consistent with previously determined product-bound structures, where CTP shares the triphosphate-binding site with UTP but adopts a distinct pyrimidine base recognition **(Fig. 6b)**. The overall conformation of the p-state is in an open state compared to RXN-state 1 **(Fig. 6b)**. These findings indicate that upon reaction completion and transition to the open state, GTP dissociates from CTPS.

**Figure 6.**
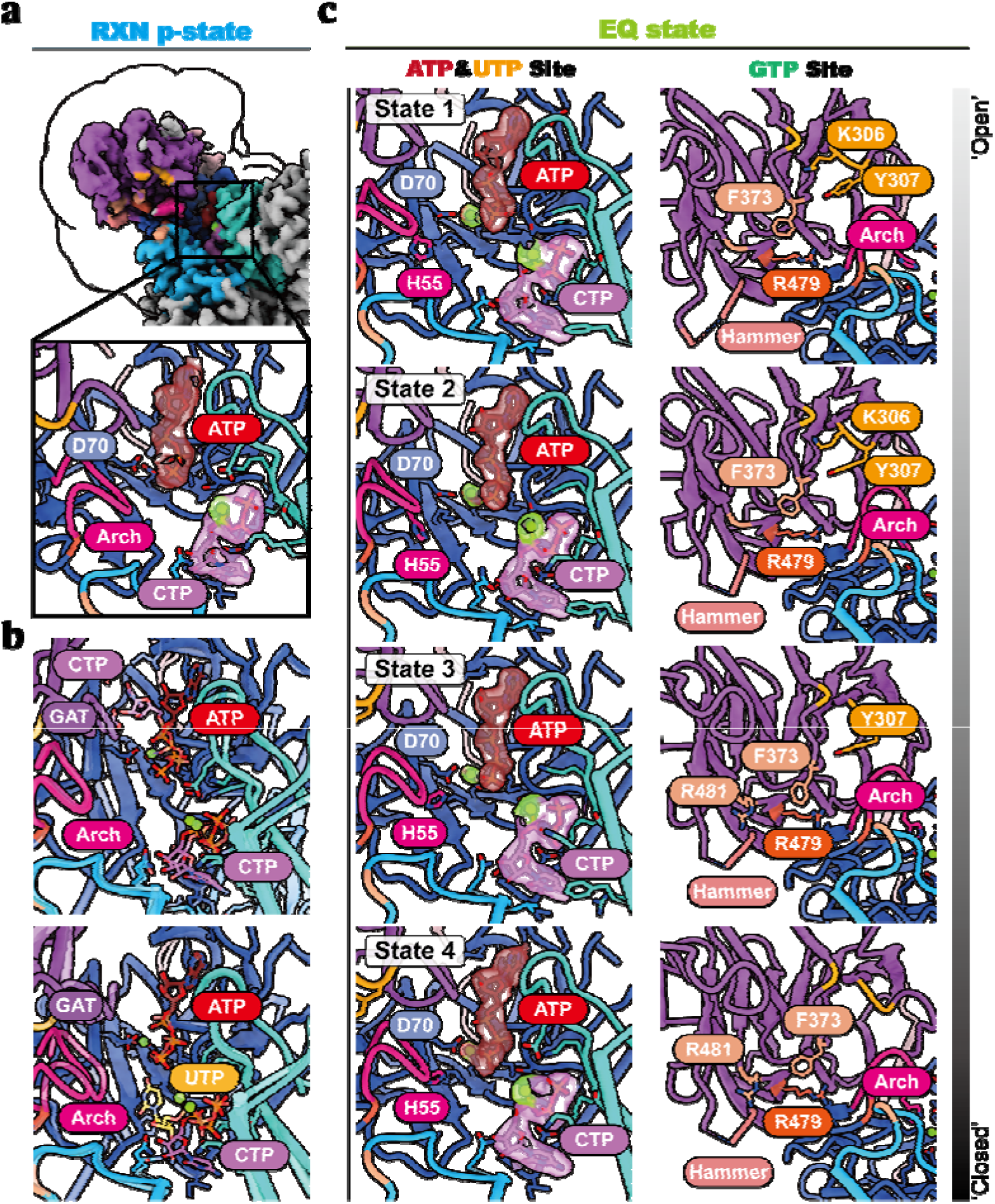
GTP unbinds post-reaction. **a**, Cryo-EM density map of the RXN p-state. A zoom-in view of the AL domain is highlighted in the boxed area. One protomer and two AL domains from adjacent protomers are shown, with different colors distinguishing each component. **b**, Comparison of ligand-binding modes in the AL domain among the RXN p-state, the previously reported CTP-inhibited state (PDB: 7PDW), and RXN state 1. The structures are aligned based on the AL domain for direct comparison. **c**, Models of all classified states from the ACP-state sample. The ligand-binding states of the AL domain and GTP-binding pocket are shown. Ligand densities are visualized as transparent surfaces.

Structures from the EQ-state dataset further support this conclusion **(Fig. 6c)**. In EQ-state 1–4, all structures show ATP and CTP co-occupying the AL domain ligand pocket, indicating an inhibited state—consistent with our enzymatic assays. No GTP binding was observed in the GAT domain, suggesting that under equilibrium conditions, GTP dissociates from CTPS.

## Discussion

### GTP cycling as a dynamic gating mechanism

Our cryo-EM analysis of 34 catalytic states reveals GTP cycling—intact binding and dissociation—serves as a molecular gatekeeper synchronising CTPS activity through three defined checkpoints (**Fig. 7**). First, GTP dissociation primes the enzyme for substrate entry by inducing an open active-site conformation. Second, 4Pi-UTP formation licenses GTP rebinding through GAT domain remodeling, coupling intermediate generation to effector recruitment. Finally, GTP-bound CTPS stabilises the ammonia transfer tunnel while coordinating catalytic residue alignment, facilitating the release of ammonia. Crucially, GTP dissociates only after CTP synthesis completes, with each catalytic round consuming precisely one GTP-binding event without hydrolysis. This cyclic coordination resolves the intricate role of GTP in CTPS catalysis, establishing effector cycling as a timing mechanism rather than energetic driver.

**Figure 7.**
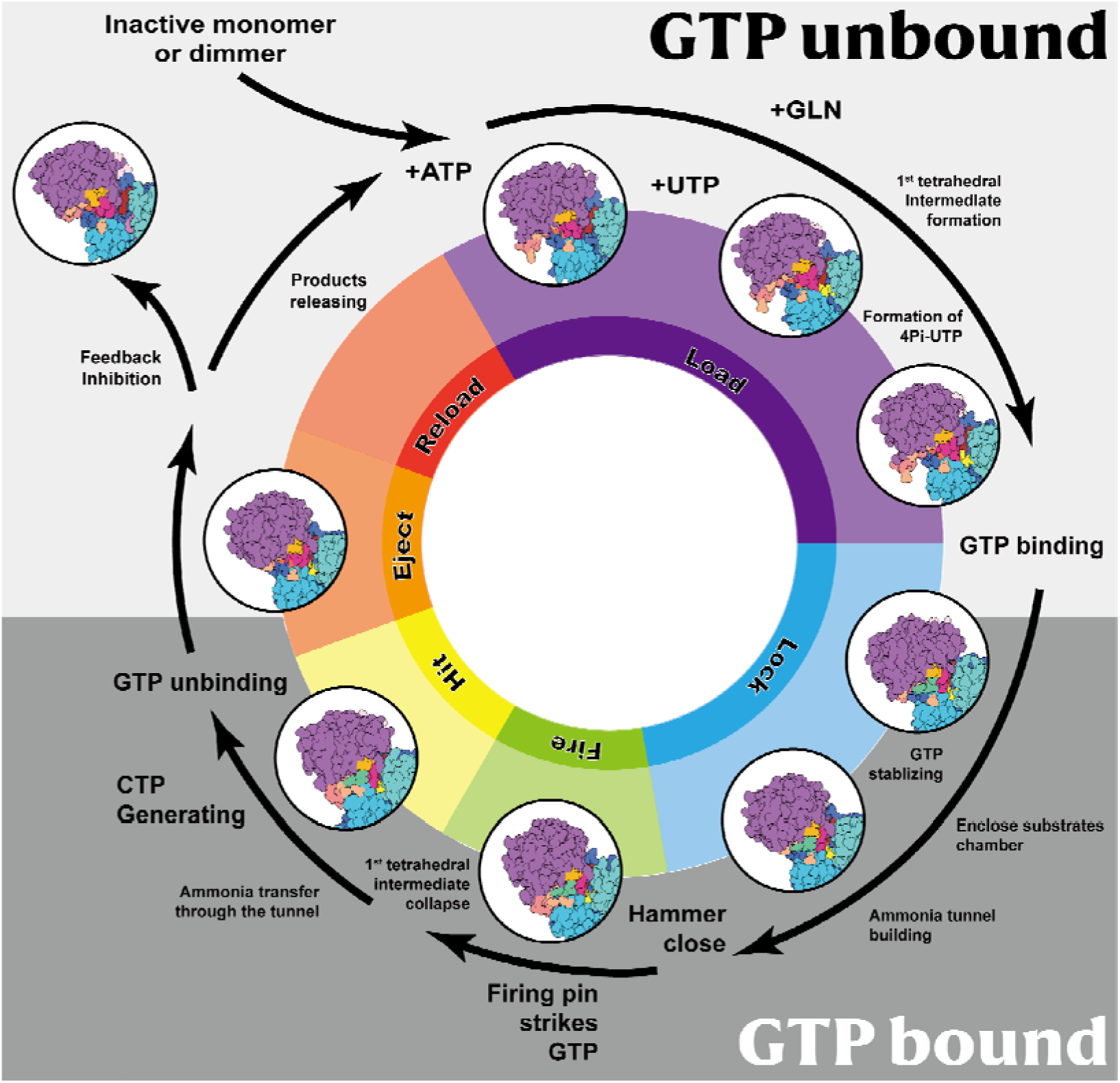
GTP cycling gates CTP synthase activation. The proposed diagram for the GTP-activated CTPS shows a single binding and dissociation event of GTP during one catalytic cycle of CTPS, where GTP binding constitutes the ammonia tunnel, facilitates GAT reaction and its subsequent dissociation allows product release.

### Evolutionary conservation of GTP-gated catalysis

The GTP-binding pocket shows remarkable conservation across archaeal, bacterial, and eukaryotic lineages, suggesting GTP cycling represents an ancient regulatory strategy. This evolutionary constraint likely reflects dual functional demands: 1) Precise coordination between cytosine production (CTP) and guanine sensing (GTP) to maintain nucleotide homeostasis; 2) Metabolic flexibility through dose-dependent regulation, where GTP levels gate CTP synthesis without irreversible effector consumption.

### Multiscale allostery in enzymatic regulation

Allostery has been hailed as the second secret of life^32,33^. CTPS exemplifies how biological systems integrate multiple allosteric strategies: 1) Conformational selection; 2) Heterotopic allostery; 3) Long range allostery and 4) Cross-talk allostery.

Our findings extend the paradigm of nucleotide-driven regulation beyond classical allostery or energy coupling, demonstrating effector cycling as a third modality where molecular binding-dissociation kinetics directly control catalytic timing. This mechanism likely operates in other nucleotide-dependent systems requiring non-destructive regulatory inputs.

## Data availability

The atomic coordinate of DON-state CTPS has been deposited in Protein Data Bank (PDB) under accession code 9UAR. The atomic coordinates of PRE-state CTPS have been deposited in PDB under accession codes 9UEY, 9UEZ, 9UF0, 9UF1, 9UF2, 9UF3, 9UF4 and 9UF5, respectively. The atomic coordinates of RXN-state CTPS have been deposited in PDB under accession codes 9V1M, 9V1N, 9V1O, 9V1P, 9V1Q, 9V1R, 9V1S, 9V1T and 9V1U, respectively. The atomic coordinates of EQ-state CTPS have been deposited in PDB under accession codes 9V1V, 9V1W, 9V1X and 9V1Y, respectively. The atomic coordinates of ACP-state CTPS have been deposited in PDB under accession codes 9V1Z, 9V20, 9V22 and 9V23, respectively. The atomic coordinates of dDON-state CTPS have been deposited in PDB under accession codes 9V2B, 9V2C, 9V2D, 9V2E, 9V2F, 9V2G, 9V2H and 9V2I, respectively. Cryo-EM map of DON-state CTPS has been deposited in the Electron Microscopy Data Bank (EMDB) under accession code EMD-63990. Cryo-EM maps of PRE-state CTPS have been deposited in the EMDB under accession codes EMD-64094, EMD-64095, EMD-64096, EMD-64097, EMD-64098, EMD-64099, EMD-64100 and EMD-64101, respectively. Cryo-EM maps of RXN-state CTPS have been deposited in the EMDB under accession codes EMD-64698, EMD-64699, EMD-64700, EMD-64701, EMD-64702, EMD-64703, EMD-64704, EMD-64705 and EMD-64706, respectively. Cryo-EM maps of EQ-state CTPS have been deposited in the EMDB under accession codes EMD-64707, EMD-64708, EMD-64709 and EMD-64710, respectively. Cryo-EM maps of ACP-state CTPS have been deposited in the EMDB under accession codes EMD-64711, EMD-64712, EMD-64714 and EMD-64715, respectively. Cryo-EM maps of dDON-state CTPS have been deposited in the EMDB under accession codes EMD-64723, EMD-64724, EMD-64725, EMD-64726, EMD-64727, EMD-64728, EMD-64729 and EMD-64730, respectively.

## Acknowledgements

We thank Suwen Zhao for her helpful discussions. EM data were collected at the ShanghaiTech Cryo-EM Imaging Facility. We thank the Molecular and Cell Biology Core Facility at the School of Life Science and Technology, ShanghaiTech University and Shanghai Frontiers Science Center for Biomacromolecules and Precision Medicine for providing technical support. This work was supported by the grants from the Ministry of Science and Technology of China (No. 2021YFA0804700), National Natural Science Foundation of China (Grant Nos. 32370744 and 32350710195), and UK Medical Research Council (Grant Nos. MC_UU_12021/3 and MC_U137788471) for grants to J. L. L.

## Contributions

C.J.G. conceptualized the study. C.J.G., Y.F.W., S.Y.G., Z.R.Z. generated the methodology. C.J.G., Y.F.W., S.Y.G., J.L.Lu, L.X., X.Z., J.Z., W.W. and Z.R.Z. performed the investigation. C.J.G., Y.F.W., S.Y.G., J.L.Lu, L.X. and Y.F. performed formal analysis. C.J.G. wrote the original draft of the manuscript. C.J.G. and Y.F. performed the visualization. C.J.G. and J.L.Liu reviewed and edited the manuscript. J.L.Liu acquired funding and provided resources. C.J.G. and J.L.Liu supervised the study.

