## Supplementary material for "GTP cycling gates CTP synthase activation": Methods

**Protein expression and purification**

Wild-type and mutant full-length CTPS proteins were expressed using previously described protocols^1,2^.

**Cryo-EM sample preparation**

Cryo-EM samples were prepared using a Vitrobot Mark IV under >90% humidity and at 4°C. Grids used were Morphous alloy film (No. M024-Au300-R12/13) and blotting was performed with TED PELLA 595 Filter Paper. Protein concentrations ranged from 1.2 to 4 μM. For inhibitor-bound conditions, the following ligand combinations were used: 1. DON-state: 2 mM UTP, 2 mM ATP, 2 mM GTP, 6 μM DON, and 10 mM MgCl₂. 2. dDON-state: 2 mM dUTP, 2 mM dATP, 2 mM dGTP, 30 μM DON, and 10 mM MgCl₂. 3. ACP-state: 2 mM UTP, 2 mM ACP, 2 mM GTP, 10 mM glutamine, and 10 mM MgCl₂

For the native ligand conditions mimicking enzymatic activity assay, samples were prepared as follows: 1. PRE-state: 2 mM ATP, 2 mM UTP, 2 mM GTP, and 10 mM MgCl₂ were incubated with protein for 3 minutes before plunge freezing. 2. RXN-state: The above mix was supplemented with 10 mM glutamine to initiate the reaction, and samples were vitrified within 3 minutes. 3. EQ-state: Samples were vitrified 15 minutes after reaction initiation, corresponding to the equilibrium state.

**Cryo-EM data collection**

All cryo-EM data were collected using a 300 kV Titan Krios G3i electron microscope equipped with a K3 camera. Automated data collection was performed using SerialEM^3^. For the DON-state and ACP-state samples, magnification was set to 22,500×, defocus range was 1.0–2.5 μm, and pixel size was 0.53 Å in super-resolution mode. The total dose was 60 e⁻/Å² distributed over 50 frames. For the dDON-state, PRE-state, RXN-state, and EQ-state samples, magnification was set to 29,000×, defocus range was 0.8–2.0 μm, and pixel size was 0.41 Å in super-resolution mode. The total dose was 59.5 e⁻/Å² distributed over 50 frames.

**Image processing and 3D reconstruction**

All data processing was carried out in CryoSPARC versions 4.6.0 and 4.6.2^4^. Raw images were motion corrected by patch motion correction and contrast transfer function (CTF) estimations were estimated by patch CTF. For the DON-state, particles were first automatically picked and cleaned by 2D classification, resulting in approximately 890k particles (1.06 Å/pixel, box size 256). 3D homogeneous refinement with D2 symmetry yielded a structure at 2.48 Å resolution. CTF refinement^5^ improved the resolution to 2.17 Å. Subsequently, the particles were polished^6^ (0.707 Å/pixel, box size 384), producing a 1.99 Å map. To improve map quality, density subtraction was performed to remove density outside the central tetramer. Ultimately, we obtained a 1.99 Å density map, which was used for local resolution estimation and local resolution-based sharpening. The sharpened density map was used for subsequent model building and visualization.

For the PRE-state, RXN-state, EQ-state, dDON-state, and ACP-state samples, we employed a broadly similar processing strategy, consisting of two main steps: (1) Homogeneous refinement of the CTPS tetramer; (2) Based on the results of homogeneous refinement, duplicate particles were removed, and the remaining particles were subjected to symmetry expansion. Using a mask focused on a single protomer, 3D classification and 3D variability analysis^7^ were performed without particle alignment to separate different conformations. Finally, state-specific density maps were obtained through homogeneous reconstruction only. The cryo-EM data collection and model refinement of various states of CTPS are listed in supplementary Table S1-S6.

For the PRE-state sample, raw data were aligned and CTF parameters estimated. A total of 5,916,521 particles were picked from 3,787 micrographs. After 2D classification and heterogeneous refinement, 461,878 particles were selected for subsequent 3D refinement. D2 symmetry was applied during consensus refinement. Based on the consensus refinement, CTF refinement and Bayesian polishing were performed to improve resolution. Finally, a 2.67 Å density map was obtained through consensus refinement. To further analyze the flexibility and dynamics of the sample, particles were subjected to symmetry expansion. Using a mask focused on a single protomer, 3D classification was performed with a resolution limit of 6 Å and 10 classes. After removing broken classes, 1,643,125 particles were used for 3D variability analysis (3DVA) with a resolution limit of 3.0 Å, 3 components, and 30 iterations. Principal component 0 was used for intermediate 3DVA display processing with a window size of 0, sorting particles into 8 non-overlapping classes. These were subsequently reconstructed to generate density maps for each state. The particle numbers and resolutions of the maps were: 67,194 (3.06 Å), 159,425 (2.98 Å), 268,077 (2.94 Å), 331,273 (2.89 Å), 315,995 (2.89 Å), 242,316 (2.87 Å), 144,921 (2.94 Å), and 64,408 (2.98 Å).

For the ACP-state sample, raw data were aligned and CTF parameters estimated. A total of 4,361,293 particles were picked from 3,669 micrographs. After 2D classification and heterogeneous refinement, 310,047 particles were selected for 3D refinement. D2 symmetry was applied during consensus refinement. Based on the consensus refinement, CTF refinement and Bayesian polishing were performed to improve resolution. A 2.71 Å density map was obtained. To further analyze sample flexibility and dynamics, particles were subjected to symmetry expansion. Using a mask focused on a single protomer, 3D classification was carried out with a resolution limit of 6 Å and 10 classes. After discarding broken classes, 1,127,358 particles were used for 3DVA with a resolution limit of 3.0 Å, 3 components, and 30 iterations. Principal component 0 was subjected to intermediate 3DVA display processing with a window size of 0, sorting particles into 4 non-overlapping classes that were reconstructed to generate density maps. The number of particles and corresponding resolutions were: 106,663 (3.09 Å), 463,586 (2.80 Å), 438,943 (2.86 Å), and 101,825 (3.09 Å).

For the dDON-state sample, raw data were aligned and CTF parameters estimated. A total of 25,405,604 particles were picked from 4,497 micrographs. After 2D classification and heterogeneous refinement, 639,063 particles were selected for 3D refinement. D2 symmetry was applied during consensus refinement. Based on the consensus refinement, CTF refinement and Bayesian polishing were used to improve the resolution. A 2.73 Å density map was obtained. To further analyze sample flexibility and dynamics, particles were subjected to symmetry expansion. Using a mask focused on a single protomer, 3D classification was performed with a resolution limit of 6 Å and 10 classes. After removing broken classes, 2,153,365 particles were used for 3DVA with a resolution limit of 3.0 Å, 3 components, and 30 iterations. Principal component 0 was subjected to intermediate 3DVA display processing with a window size of 0, sorting particles into 8 non-overlapping classes that were subsequently reconstructed to generate density maps. The particle numbers and resolutions were: 97,100 (3.15 Å), 264,542 (2.98 Å), 426,024 (3.00 Å), 423,774 (2.99 Å), 339,502 (3.00 Å), 272,662 (2.96 Å), 182,940 (2.93 Å), and 82,959 (3.01 Å).

For the RXN-state sample, raw data were aligned and CTF parameters estimated. A total of 5,527,860 particles were picked from 8,134 micrographs. After 2D classification and heterogeneous refinement, 714,929 particles were selected for subsequent 3D refinement. D2 symmetry was applied during consensus refinement. Based on the consensus refinement, CTF refinement and Bayesian polishing were performed to improve resolution. A 2.76 Å density map was obtained through consensus refinement. To further investigate the flexibility and dynamics of the sample, particles were subjected to symmetry expansion. Using a mask focused on a single protomer, 3D classification was performed with a resolution limit of 6 Å and 10 classes. One of the classes showed CTP binding in the AL domain; this class was processed separately, and a further subclassification followed by reconstruction using 33,365 particles yielded a 3.25 Å structure of the p-state. After discarding broken classes, 1,643,125 particles were used for 3D variability analysis (3DVA) with a resolution limit of 3.0 Å, 3 components, and 30 iterations. Principal component 0 was subjected to intermediate 3DVA display processing with a window size of 0, which sorted particles into 8 non-overlapping classes. These classes were reconstructed to generate density maps of the corresponding states. The particle numbers and resolutions for each map were: 79,751 (3.20 Å), 202,371 (3.05 Å), 377,549 (2.98 Å), 492,744 (2.98 Å), 447,630 (3.00 Å), 308,790 (2.97 Å), 173,920 (3.00 Å), and 78,103 (3.08 Å).

For the EQ-state sample, raw data were aligned and CTF parameters estimated. A total of 405,455 particles were picked from 5,856 micrographs. After 2D classification and heterogeneous refinement, 358,641 particles were selected for 3D refinement. D2 symmetry was applied during consensus refinement. Based on the consensus refinement, CTF refinement was performed to improve resolution. A 2.78 Å density map was obtained. To further analyze sample flexibility and dynamics, particles were subjected to symmetry expansion. Using a mask focused on a single protomer, 3D classification was performed with a resolution limit of 6 Å and 10 classes. After discarding broken classes, 1,273,536 particles were used for 3DVA with a resolution limit of 3.0 Å, 3 components, and 30 iterations. Principal component 0 was subjected to intermediate 3DVA display processing with a window size of 0, which sorted particles into 4 non-overlapping classes. These were subsequently reconstructed to generate density maps. The number of particles and corresponding resolutions were: 128,100 (3.14 Å), 497,433 (2.96 Å), 501,894 (2.95 Å), and 128,048 (3.10 Å).

Local resolution estimates were made for all maps using cryoSPARC with an FSC cut-off of 0.143. Local filtering and DeepEMhancer^8^ were used for map sharpening.

**Model building**

For the DON-state dataset, model building began using PDB entry 7DPT as the initial model. This model was manually adjusted and refined against the local resolution-sharpened map using Phenix^9,10^. Solvent water molecules were then added automatically in Coot^11^ and manually curated. The final model was obtained after multiple rounds of real-space refinement in Phenix.

For the PRE-state, ACP-state, RXN-state, EQ-state, and dDON-state datasets, an identical strategy was employed: (1) Initial models (7DPW for the ‘open’ states and 7DPT for the ‘closed’ states) were rigid-body fit into the DeepEM-sharpened maps. (2) Initial real-space refinement was performed in Phenix. The map-model simulation in ISOLDE^12^ was used to optimize models. (3) NTP and dNTP ligands were manually added to the models, followed by real-space refinement against the local resolution-sharpened maps in Phenix to generate the final structures. For the dDON-state dataset, where ligand density was observed in the GAT domain, the GAT domain from the DON-state was fit in, DON was modeled into the density, and refined using Phenix.

For glutamine-containing samples, due to the catalytic turnover of glutamine by the GAT domain and resolution limitations, no ligands were modeled in the GAT domain. Glutamine (GLN) was instead modeled as a bound ligand by rigid-body fitting into the density, solely for ligand density visualization purposes.

**Tunnel modeling.**

CAVER Analyst^13,14^ was used to analyze and model the ammonia tunnel of DON-state CTPS. The starting point for tunnel detection was defined by manually selecting amino acids surrounding Cys399. Tunnel visualization was performed in ChimeraX^15^.

**Enzyme assay for CTPS**

The activity of the CTPS GAT domain was determined using the previously described two-step Glutaminase protocol based on L-Glutamate Dehydrogenase (GDH)^16,17^. CTP production was monitored by measuring absorbance at 291 nm^18^. A typical reaction was set up by incubating 2 μM purified CTPS protein in a 96-well plate containing 2 mM ATP, 2 mM UTP, 0.2 mM GTP, 10 mM MgCl₂, and buffer (50 mM Tris-HCl, pH 8.0, 150 mM NaCl) in a total volume of 200 μL. After a 15-minute incubation at 37°C, the reaction was initiated by adding 10 mM glutamine prewarmed to 37°C. Absorbance measurements were recorded at 10-second intervals using a SpectraMax i3 plate reader.

**Sequence conservation analysis**
All sequences used for multiple sequence alignment were obtained from the UniProt database^19^. Sequences were selected from different phyla based on the UniProt taxonomy, using EC number 6.3.4.2 as the search criterion. Multiple sequence alignments were performed using MAFFT^20^ on the online platform^21^ <https://toolkit.tuebingen.mpg.de/>. Visualization of sequence conservation was carried out using ESPript3^22^ (<https://espript.ibcp.fr/ESPript/ESPript/>) and WebLogo3^23^ (<https://weblogo.threeplusone.com/create.cgi>).

1	Zhou, X. *et al.* Drosophila CTP synthase can form distinct substrate-and product-bound filaments. *Journal of Genetics and Genomics* **46**, 537-545 (2019).

2	Zhou, X. *et al.* Structural basis for ligand binding modes of CTP synthase. *P Natl Acad Sci USA* **118** (2021). <https://doi.org:ARTN> e2026621118

10.1073/pnas.2026621118

3	Mastronarde, D. N. Automated electron microscope tomography using robust prediction of specimen movements. *J Struct Biol* **152**, 36-51 (2005). <https://doi.org:10.1016/j.jsb.2005.07.007>

4	Punjani, A., Rubinstein, J. L., Fleet, D. J. & Brubaker, M. A. cryoSPARC: algorithms for rapid unsupervised cryo-EM structure determination. *Nat Methods* **14**, 290-296 (2017). <https://doi.org:10.1038/nmeth.4169>

5	Zivanov, J., Nakane, T. & Scheres, S. H. W. Estimation of high-order aberrations and anisotropic magnification from cryo-EM data sets in RELION-3.1. *IUCrJ* **7**, 253-267 (2020). <https://doi.org:10.1107/S2052252520000081>

6	Zivanov, J., Nakane, T. & Scheres, S. H. W. A Bayesian approach to beam-induced motion correction in cryo-EM single-particle analysis. *IUCrJ* **6**, 5-17 (2019). <https://doi.org:10.1107/S205225251801463X>

7	Punjani, A. & Fleet, D. J. 3D variability analysis: Resolving continuous flexibility and discrete heterogeneity from single particle cryo-EM. *J Struct Biol* **213**, 107702 (2021). <https://doi.org:10.1016/j.jsb.2021.107702>

8	Sanchez-Garcia, R. *et al.* DeepEMhancer: a deep learning solution for cryo-EM volume post-processing. *Commun Biol* **4**, 874 (2021). <https://doi.org:10.1038/s42003-021-02399-1>

9	Adams, P. D. *et al.* PHENIX: a comprehensive Python-based system for macromolecular structure solution. *Acta Crystallogr D Biol Crystallogr* **66**, 213-221 (2010). <https://doi.org:10.1107/s0907444909052925>

10	Afonine, P. V. *et al.* Real-space refinement in PHENIX for cryo-EM and crystallography. *Acta Crystallogr D Struct Biol* **74**, 531-544 (2018). <https://doi.org:10.1107/s2059798318006551>

11	Emsley, P., Lohkamp, B., Scott, W. G. & Cowtan, K. Features and development of Coot. *Acta Crystallogr D Biol Crystallogr* **66**, 486-501 (2010). <https://doi.org:10.1107/S0907444910007493>

12	Croll, T. I. ISOLDE: a physically realistic environment for model building into low-resolution electron-density maps. *Acta Crystallogr D Struct Biol* **74**, 519-530 (2018). <https://doi.org:10.1107/S2059798318002425>

13	Jurcik, A. *et al.* CAVER Analyst 2.0: analysis and visualization of channels and tunnels in protein structures and molecular dynamics trajectories. *Bioinformatics* **34**, 3586-3588 (2018). <https://doi.org:10.1093/bioinformatics/bty386>

14	Kozlikova, B. *et al.* CAVER Analyst 1.0: graphic tool for interactive visualization and analysis of tunnels and channels in protein structures. *Bioinformatics* **30**, 2684-2685 (2014). <https://doi.org:10.1093/bioinformatics/btu364>

15	Goddard, T. D. *et al.* UCSF ChimeraX: Meeting modern challenges in visualization and analysis. *Protein Sci* **27**, 14-25 (2018). <https://doi.org:10.1002/pro.3235>

16	Willemoës, M. & Sigurskjold, B. W. Steady-state kinetics of the glutaminase reaction of CTP synthase from Lactococcus lactis. The role of the allosteric activator GTP incoupling between glutamine hydrolysis and CTP synthesis. *Eur J Biochem* **269**, 4772-4779 (2002). <https://doi.org:10.1046/j.1432-1033.2002.03175.x>

17	MacDonnell, J. E., Lunn, F. A. & Bearne, S. L. Inhibition of E. coli CTP synthase by the "positive" allosteric effector GTP. *Bba-Proteins Proteom* **1699**, 213-220 (2004). <https://doi.org:10.1016/j.bbapap.2004.03.002>

18	Levitzki, A. & Koshland, D. E. Cytidine triphosphate synthetase. Covalent intermediates and mechanisms of action. *Biochemistry-Us* **10**, 3365-3371 (1971). <https://doi.org:10.1021/bi00794a008>

19	UniProt, C. UniProt: the Universal Protein Knowledgebase in 2025. *Nucleic Acids Res* **53**, D609-D617 (2025). <https://doi.org:10.1093/nar/gkae1010>

20	Katoh, K., Kuma, K.-i., Toh, H. & Miyata, T. MAFFT version 5: improvement in accuracy of multiple sequence alignment. *Nucleic Acids Res* **33**, 511-518 (2005). <https://doi.org:10.1093/nar/gki198>

21	Gabler, F. *et al.* Protein Sequence Analysis Using the MPI Bioinformatics Toolkit. *Curr Protoc Bioinformatics* **72**, e108 (2020). <https://doi.org:10.1002/cpbi.108>

22	Robert, X. & Gouet, P. Deciphering key features in protein structures with the new ENDscript server. *Nucleic Acids Res* **42**, W320-324 (2014). <https://doi.org:10.1093/nar/gku316>

23	Crooks, G. E., Hon, G., Chandonia, J. M. & Brenner, S. E. WebLogo: a sequence logo generator. *Genome Res* **14**, 1188-1190 (2004). <https://doi.org:10.1101/gr.849004>
