## Supplemental Figures S1-S11 and legends for "GTP cycling gates CTP synthase activation"

#### Supplementary Figures S1-S11 & Legends

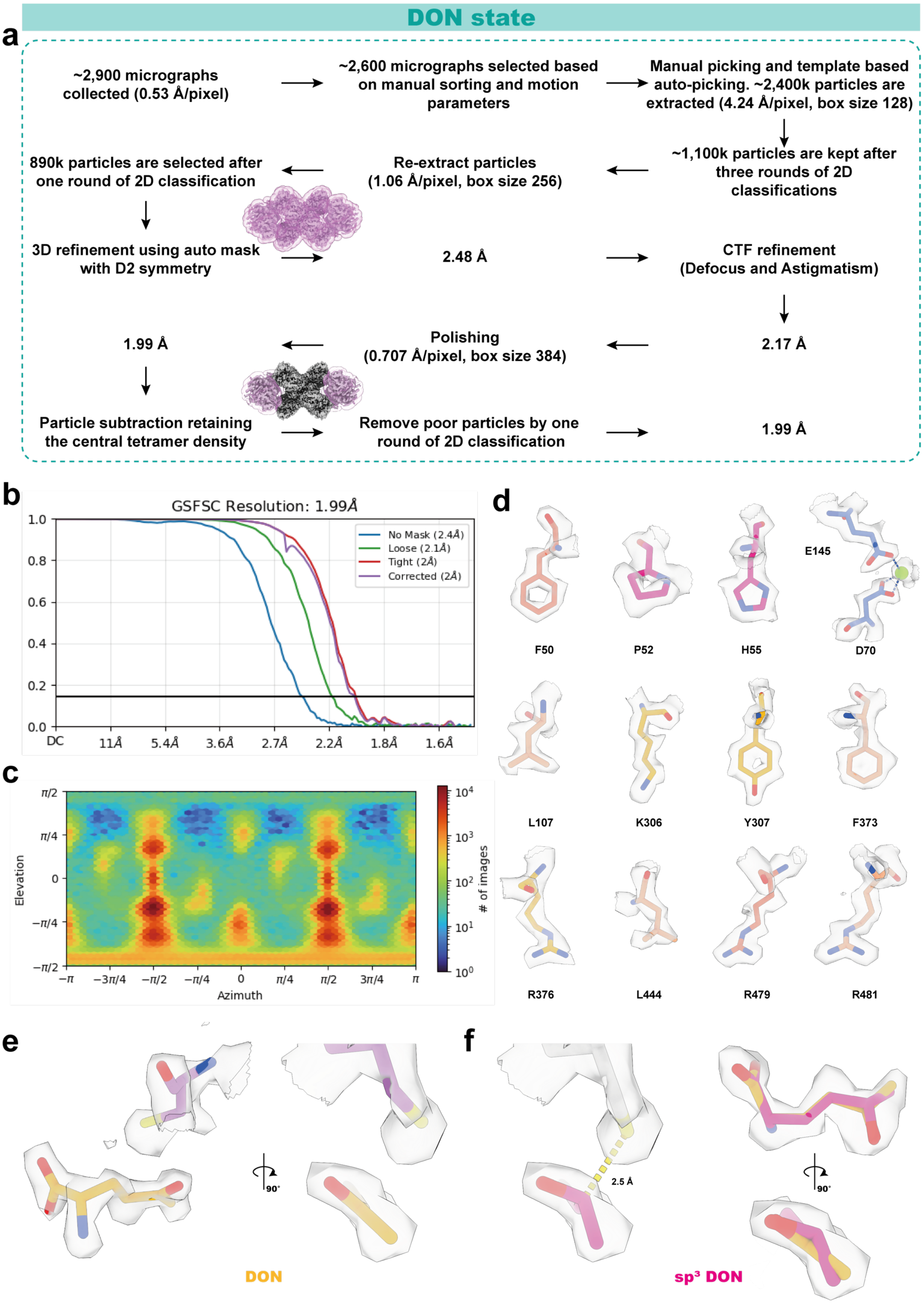
DON state
a
~2,900 micrographs collected (0.53 Å/pixel)
~2,600 micrographs selected based on manual sorting and motion parameters
Manual picking and template based auto-picking. ~2,400k particles are extracted (4.24 Å/pixel, box size 128)
~1,100k particles are kept after three rounds of 2D classifications
Re-extract particles (1.06 Å/pixel, box size 256)
890k particles are selected after one round of 2D classification
3D refinement using auto mask with D2 symmetry
2.48 Å
CTF refinement (Defocus and Astigmatism)
2.17 Å
Polishing (0.707 Å/pixel, box size 384)
1.99 Å
Particle subtraction retaining the central tetramer density
Remove poor particles by one round of 2D classification
1.99 Å
b
GSFSC Resolution: 1.99Å
No Mask (2.4Å)
Loose (2.1Å)
Tight (2Å)
Corrected (2Å)
DC
11Å
5.4Å
3.6Å
2.7Å
2.2Å
1.8Å
1.6Å
c
Elevation
Azimuth
### of images
d
F50
P52
H55
E145
D70
L107
K306
Y307
F373
R376
L444
R479
R481
e
DON
f
2.5 Å
sp³ DON

**Figure S1 | DON-state data processing.**

a, The cryo-EM image processing workflow for DON-state sample.

b, Fourier shell correlation (FSC) curve between the two half-maps obtained by the final 3D refinement.

c, Angular distribution plot for the final 3D refinement.

d, Map details for the ligand interacting amino acids.

e, Density and model of the ligand DON and catalytic amino acid cys399.

f, Fitting of the $sp^3$-hybridized DON model to the electron density. Right panel: Comparison between models of DON in $sp^2$ and $sp^3$ hybridization states.

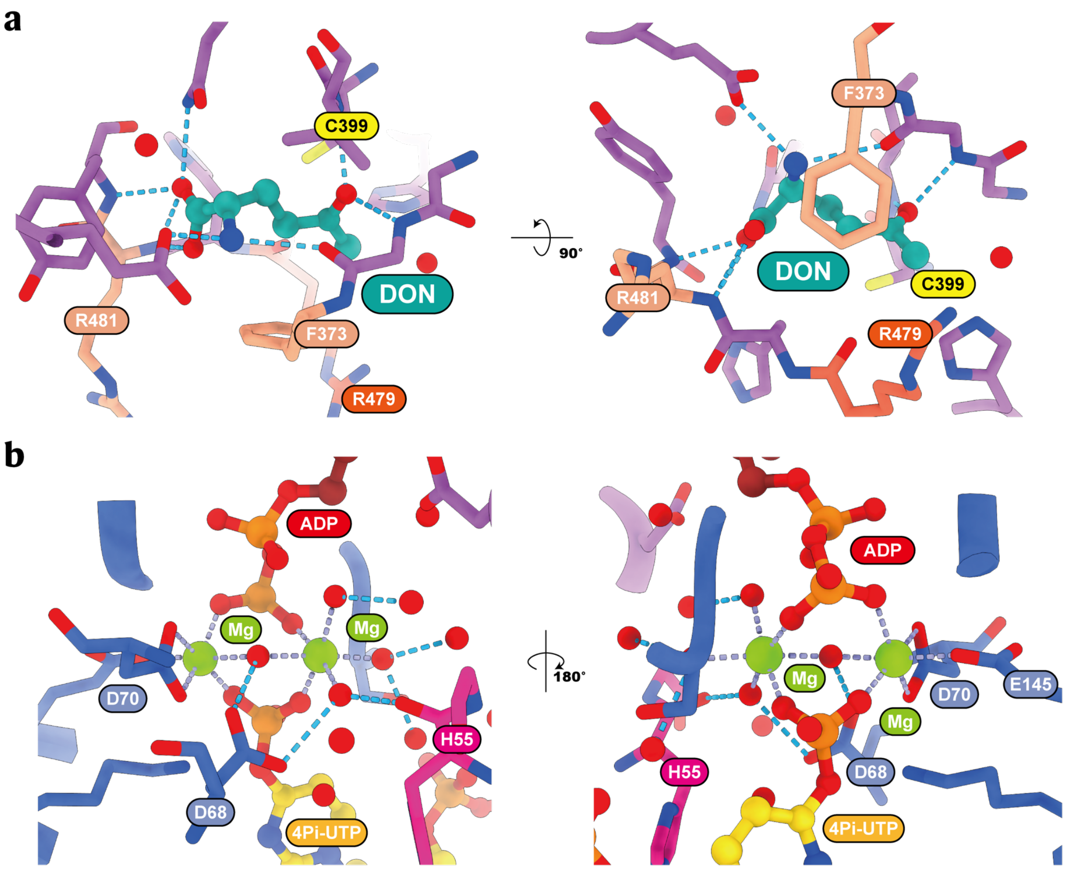


**Figure S2 | Structural details of ligand binding in DON-state.**

a, The binding mode of DON.

b, Interactions of magnesium ion with ADP, 4Pi-UTP, and solvent. The blue dashed lines represent hydrogen bonds, calculated using ChimeraX.

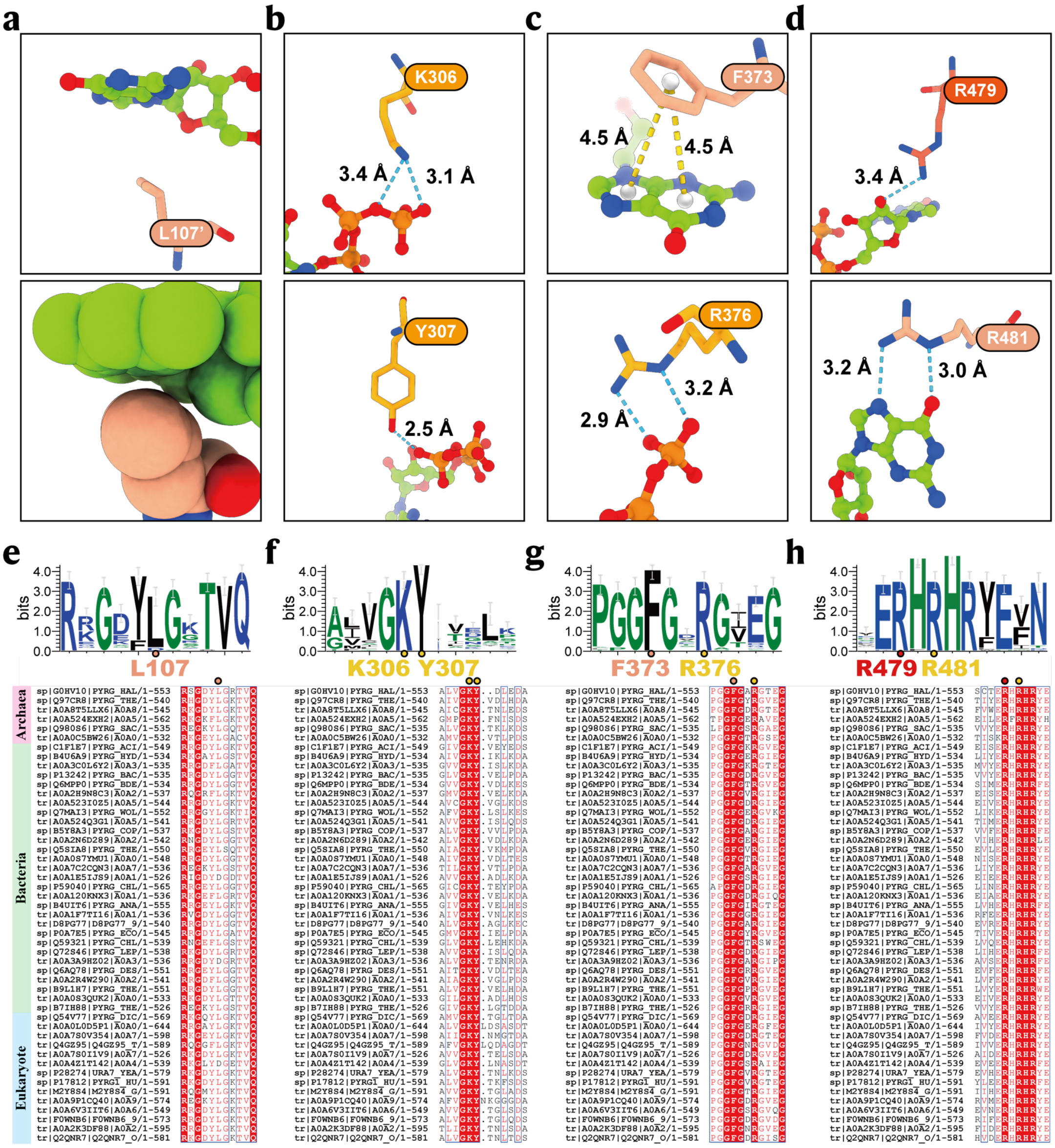


### Figure S3 | The conservativity of GTP-interacting amino acids.

a-d, Interactions of different CTPS amino acids with GTP. The lower panel of a shows the van der Waals radii of atoms in sphere style.

e-h, Conservation analysis of GTP-interacting amino acids across all three domains of life. Sequences were selected based on different phyla in the UniProt taxonomy.

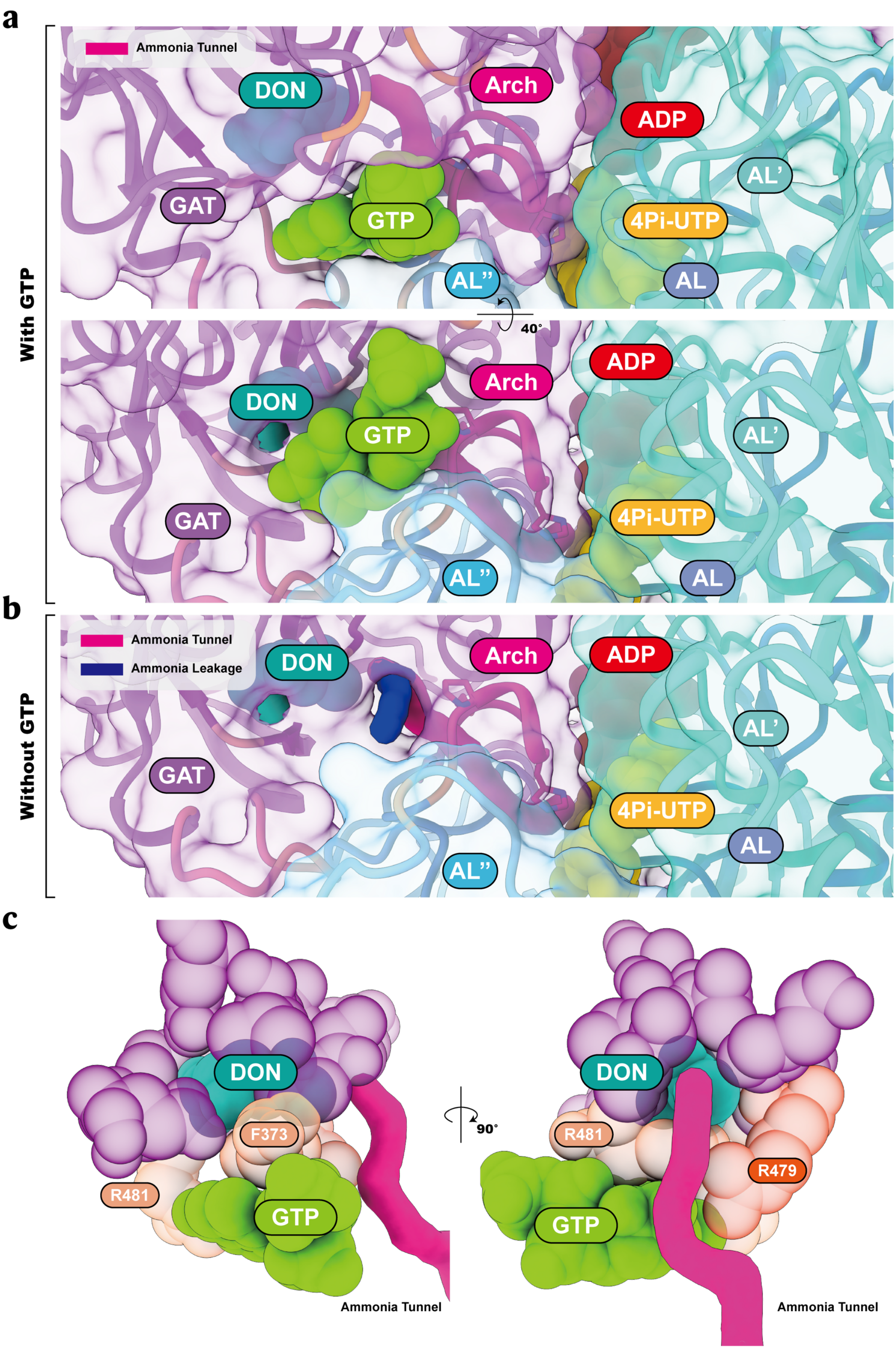
a
With GTP
Ammonia Tunnel
DON
Arch
ADP
AL'
GAT
GTP
4Pi-UTP
AL''
AL
40°
ADP
Arch
DON
GTP
AL'
GAT
4Pi-UTP
AL''
AL
b
Without GTP
Ammonia Tunnel
Ammonia Leakage
DON
Arch
ADP
AL'
GAT
4Pi-UTP
AL
AL''
c
DON
F373
R481
GTP
Ammonia Tunnel
90°
DON
R481
R479
GTP
Ammonia Tunnel

36
37

**Figure S4 | GTP constitutes the ammonia tunnel and seals the GAT pocket.**

a, Model of the ammonia channel in the DON-state. The central axis of the ammonia channel is shown as a magenta surface.

b, In the DON-state model, the absence of GTP leads to leakage of the ammonia channel.

c, GTP seals the reaction chamber of the GAT domain. Atoms are displayed in sphere style in ChimeraX showing the van der Waals (VDW) radii.

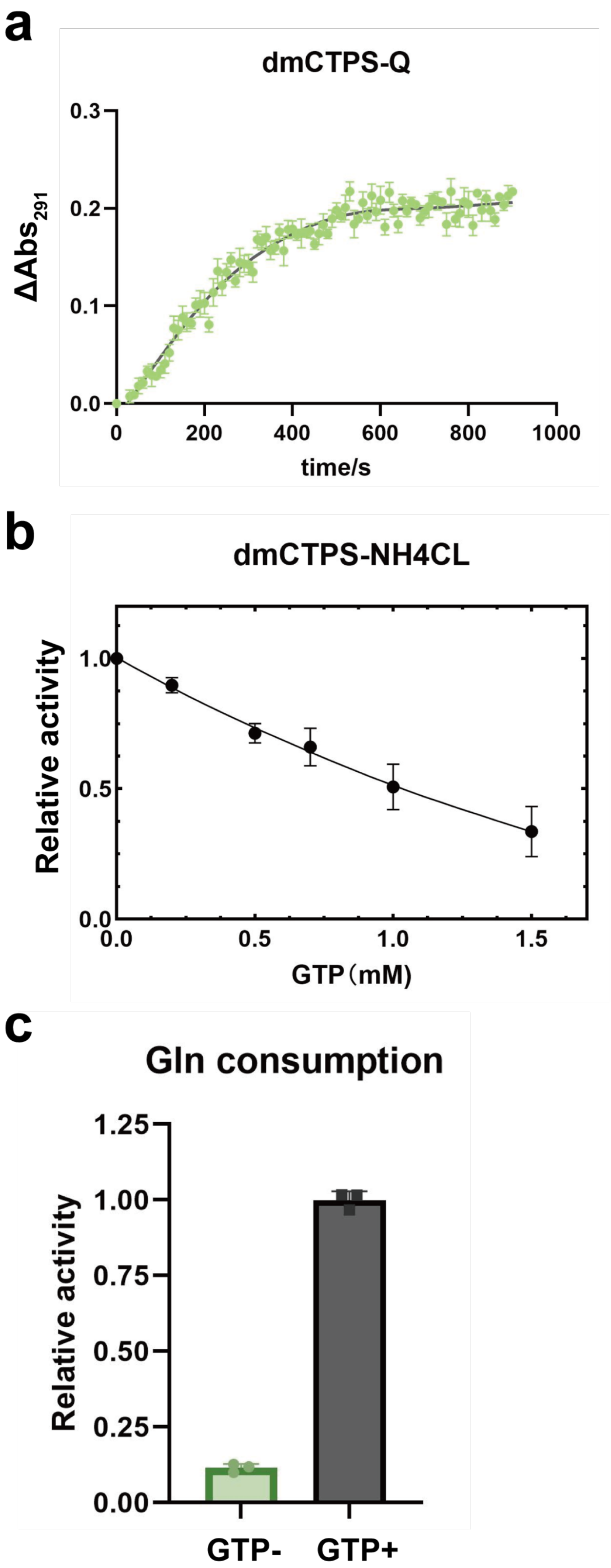
a
dmCTPS-Q
ΔAbs291
0.3
0.2
0.1
0.0
0
200
400
600
800
1000
time/s
b
dmCTPS-NH4CL
Relative activity
1.0
0.5
0.0
0.0
0.5
1.0
1.5
GTP（mM)
c
Gln consumption
Relative activity
1.25
1.00
0.75
0.50
0.25
0.00
GTP-
GTP+

**Figure S5 | Enzyme activity assays under various conditions.**

a, Change in absorbance at 291 nm after the reaction was initiated by adding 10 mM glutamine to the reaction mixture (2 μM CTPS, 2 mM ATP, UTP, GTP, 10 mM $MgCl_2$ pre-incubation).

b, Effects of different GTP concentrations (0-1.5 mM) on $NH_3$-dependent dmCTPS activity (2 μM CTPS, 2 mM ATP, UTP, 10 mM $MgCl_2$).

c, Activation of the GAT domain by GTP. The reaction was initiated with 10 mM glutamine (ATP 1mM, UTP 1mM, GTP 0 or 0.4 mM, $NADP^+$ 2mM, GDH 0.5U, CTPS 2uM 10 mM $MgCl_2$).

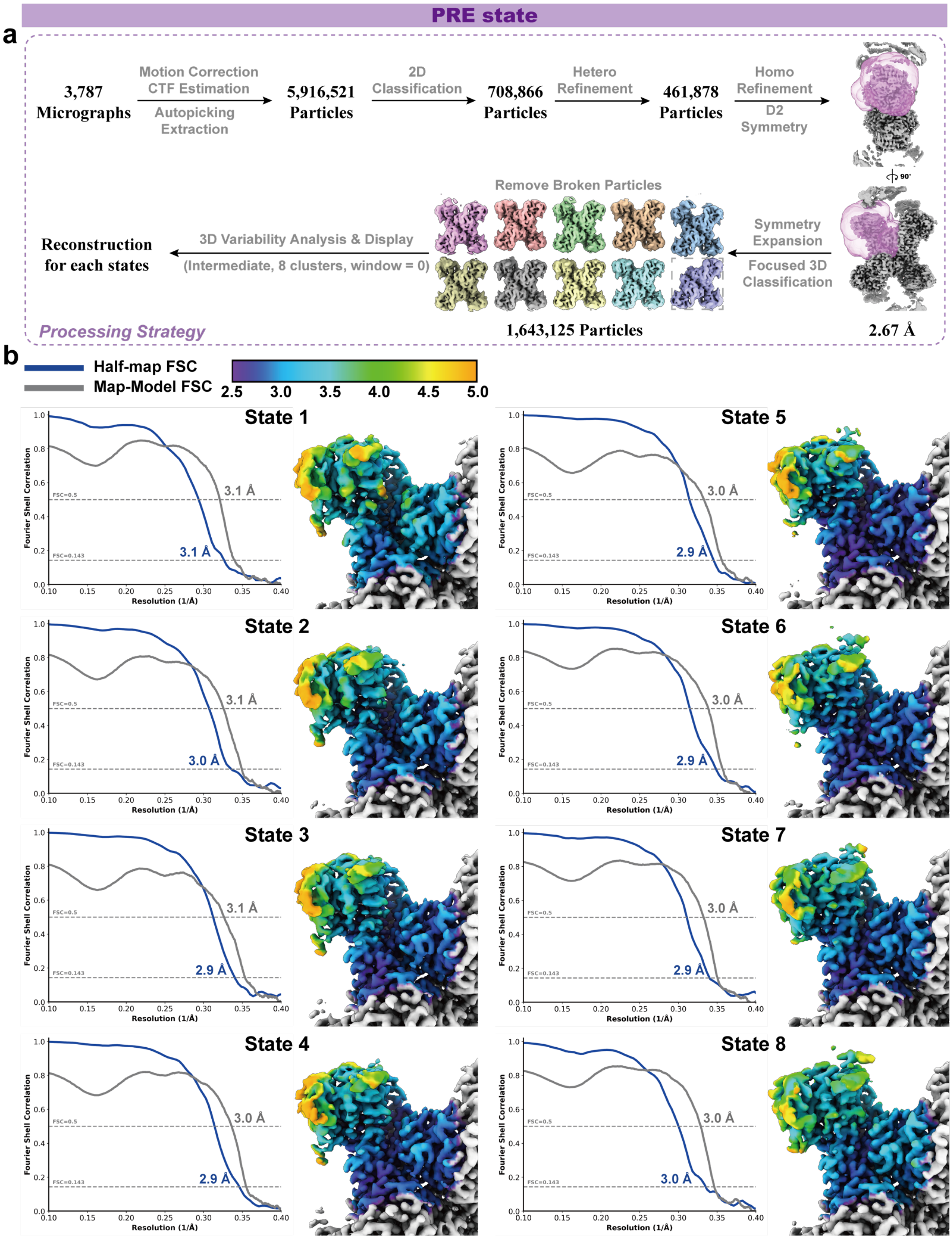


**Figure S6 | PRE-state data processing.**

a, The image processing strategy for PRE state sample.

b, Forrier shell correlation and local resolution distribution of the PRE state structures.

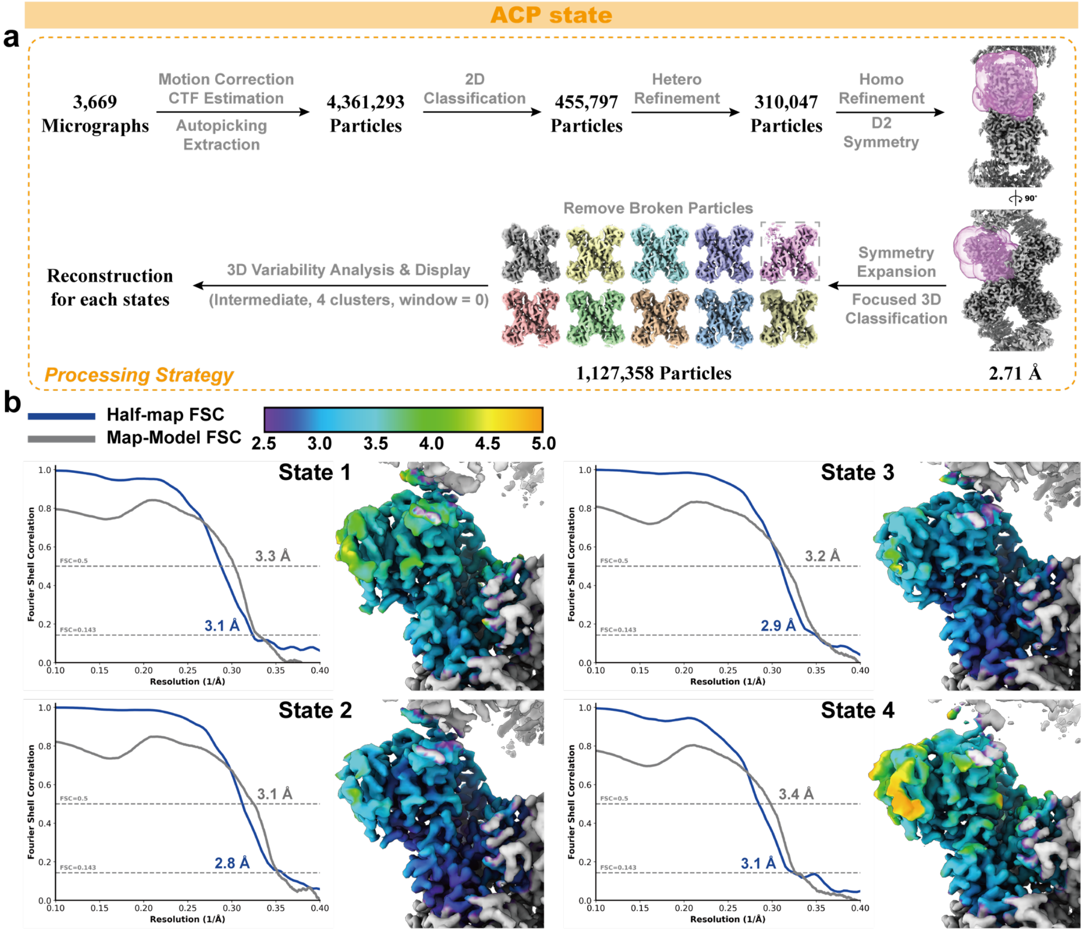


### Figure S7 | ACP-state data processing.

a, The image processing strategy for ACP state sample.

b, Forrier shell correlation and local resolution distribution of the ACP state structures.

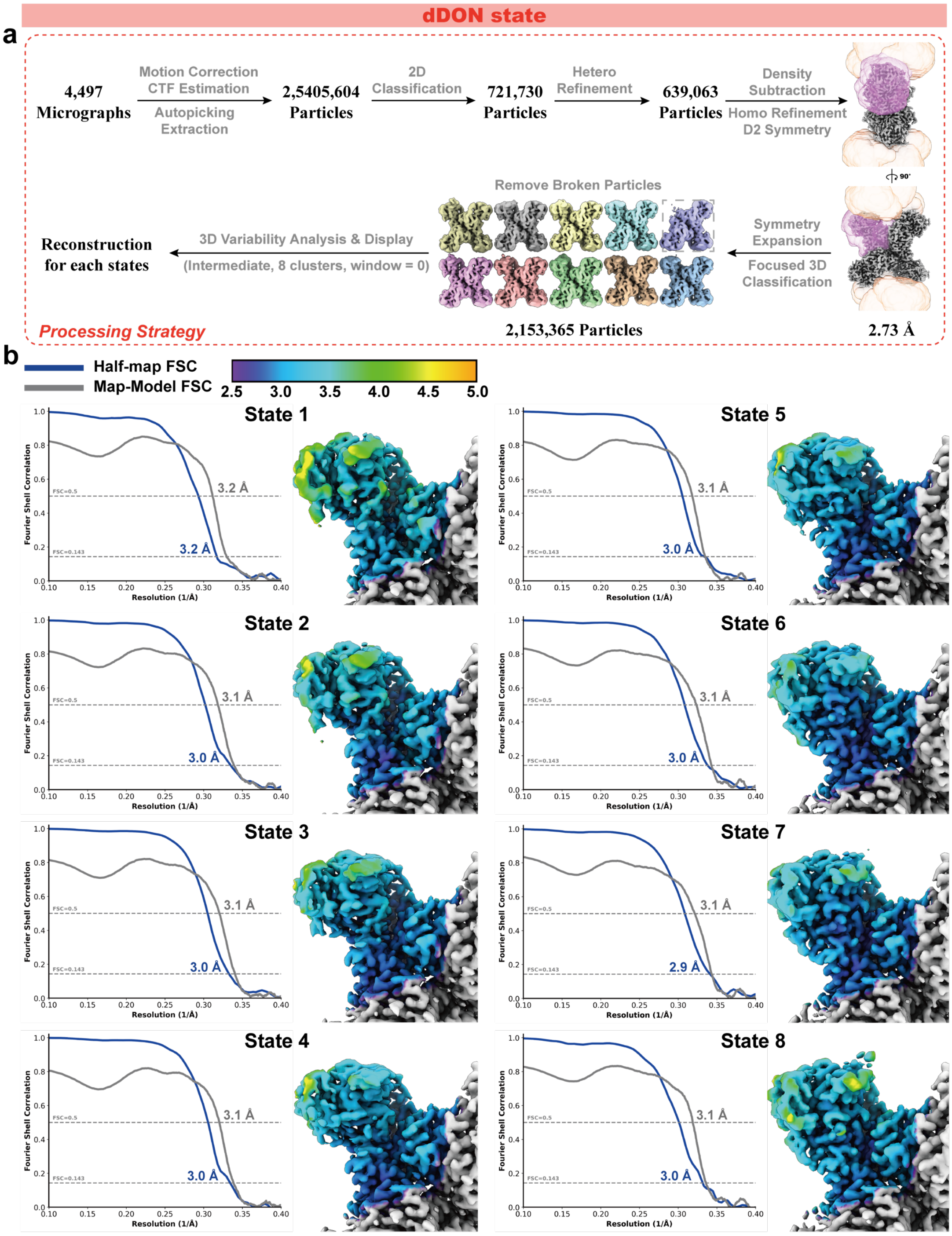


**Figure S8 | dDON-state data processing.**

a, The image processing strategy for dDON state sample.

b, Forrier shell correlation and local resolution distribution of the dDON state structures.

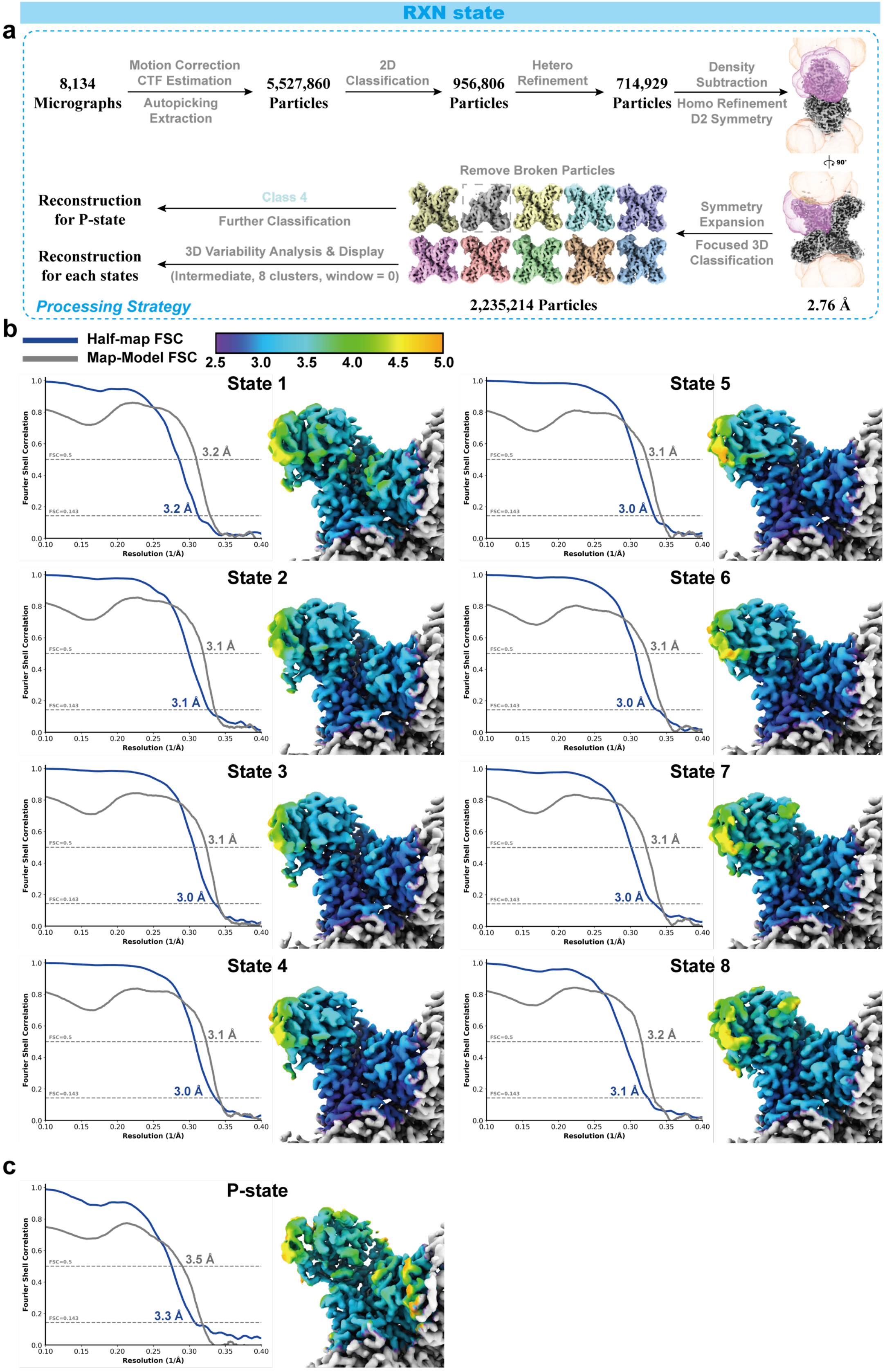
RXN state
a
8,134 Micrographs
Motion Correction
CTF Estimation
Autopicking
Extraction
5,527,860 Particles
2D Classification
956,806 Particles
Hetero Refinement
714,929 Particles
Density Subtraction
Homo Refinement
D2 Symmetry
90°
Symmetry Expansion
Focused 3D Classification
Remove Broken Particles
Class 4
Further Classification
Reconstruction for P-state
3D Variability Analysis & Display
(Intermediate, 8 clusters, window = 0)
Reconstruction for each states
Processing Strategy
2,235,214 Particles
2.76 Å
b
Half-map FSC
Map-Model FSC
2.5
3.0
3.5
4.0
4.5
5.0
State 1
3.2 Å
3.2 Å
State 2
3.1 Å
3.1 Å
State 3
3.1 Å
3.0 Å
State 4
3.1 Å
3.0 Å
State 5
3.1 Å
3.0 Å
State 6
3.1 Å
3.0 Å
State 7
3.1 Å
3.0 Å
State 8
3.2 Å
3.1 Å
Fourier Shell Correlation
Resolution (1/Å)
FSC=0.5
FSC=0.143
c
P-state
3.5 Å
3.3 Å

**Figure S9 | RXN-state data processing.**

a, The image processing strategy for RXN state sample.

b, Forrier shell correlation and local resolution distribution of the RXN state 1-8 structures.

c, Forrier shell correlation and local resolution distribution of the RXN p-state structure.

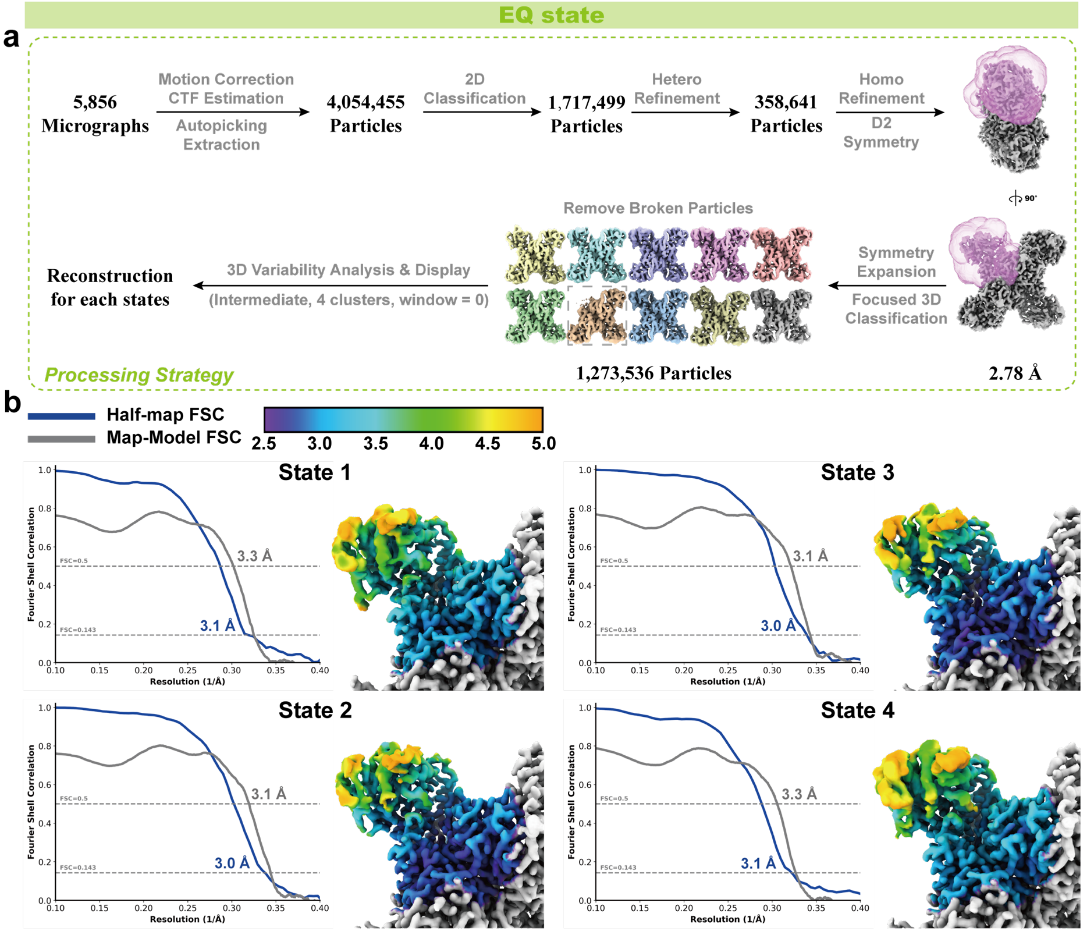


**Figure S10 | EQ-state data processing.**

a, The image processing strategy for EQ state sample.

b, Forrier shell correlation and local resolution distribution of the EQ state structures.

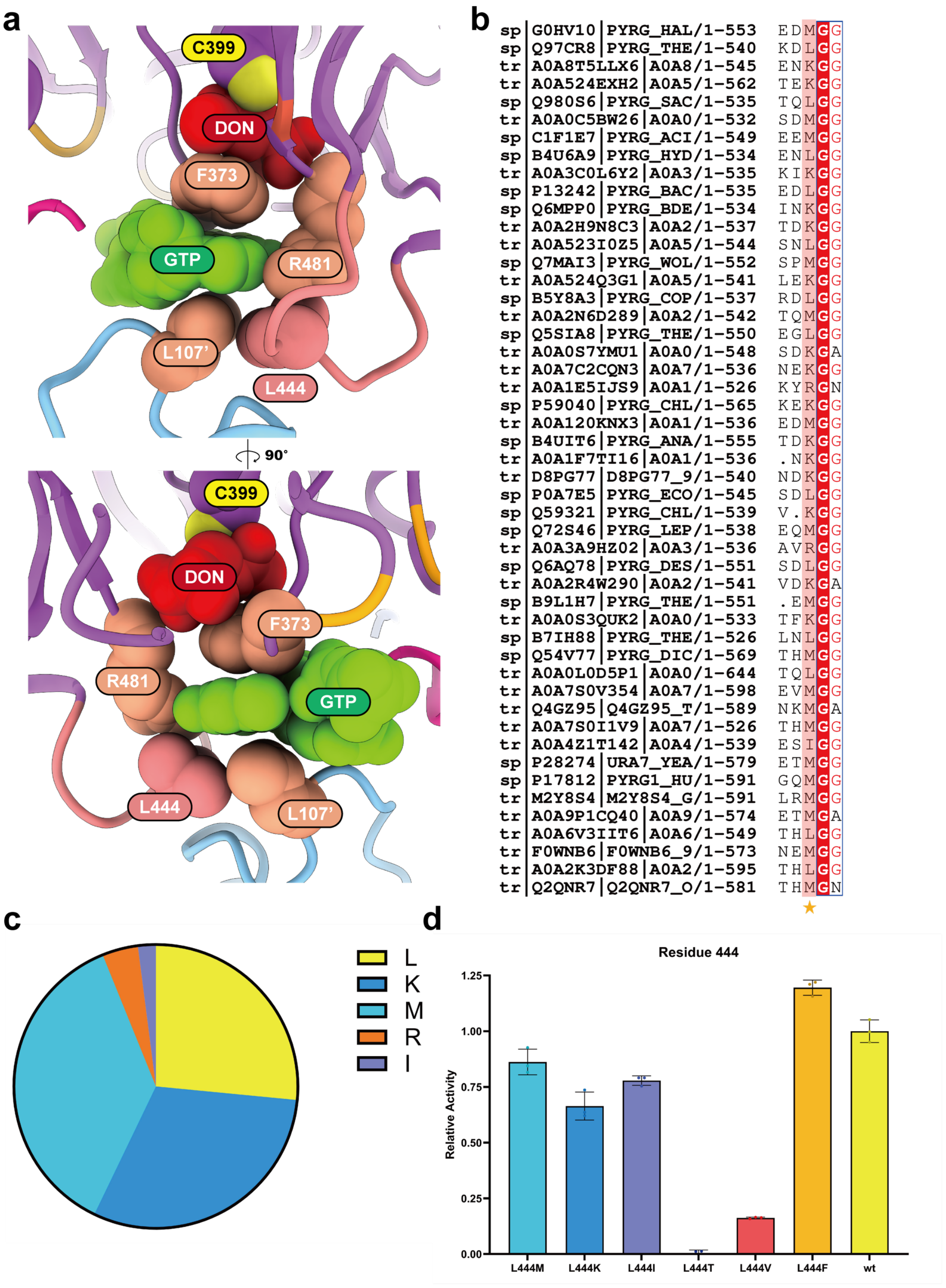


**Figure S11 | Analysis of the impact of L444 on CTPS activity.**

a, Interaction of L444 with GTP in the DON-state.

b, Multiple sequence alignment reveals the evolutionary diversity of L444.

c, Statistics on the occurrence of different amino acids at position 444.

d, Impact of residue 444 mutations on CTPS activity.
