## Supplemental Videos S1-S13 for "GTP cycling gates CTP synthase activation": Supplementary Videos.docx

**Supplementary Video Legends**

***Supplementary Video 1* | Visualization of chemical details for GTP recognition in the 2.0 Å structure.**

***Supplementary Video 2* | Visualization of chemical details for intermediate 4Pi-UTP stabilization in the 2.0 Å structure.**

***Supplementary Video 3* | Visualization of the relative spatial relationship between GTP and GAT ligands.** DON is colored in yellow, and the CTPS is displayed as a purple surface.

***Supplementary Video 4* | Video summarizing the density maps from the PRE state dataset.**

***Supplementary Video 5* | Video summarizing the structures from the PRE state dataset.**

***Supplementary Video 6* | Video summarizing the density maps from the RXN state dataset.**

***Supplementary Video 7* | Video summarizing the structures from the RXN state dataset.**

***Supplementary Video 8* | Video summarizing the density maps from the EQ state dataset.**

***Supplementary Video 9* | Video summarizing the structures from the EQ state dataset.**

***Supplementary Video 10* | Video summarizing the density maps from the ACP state dataset.**

***Supplementary Video 11* | Video summarizing the structures from the ACP state dataset.**

***Supplementary Video 12* | Video summarizing the density maps from the dDON state dataset.**

***Supplementary Video 13* | Video summarizing the structures from the dDON state dataset.**
